# Physics-based nucleosome-resolution modeling of epigenetic-driven chromatin domain dynamics

**DOI:** 10.1101/2025.08.04.668597

**Authors:** Chenyang Gu, Shoji Takada, Giovanni B. Brandani

## Abstract

Chromatin spatially organizes the Eukaryotic genome to support key cellular processes such as gene regulation, but the interplay between epigenetics, chromatin structure, and function is still poorly understood. We propose a nucleosome-resolution coarse-grained model that captures the essential features of chromatin organization over multiple scales: from nucleosome dynamics to chromatin fiber folding, and liquid-liquid phase separation. The model describes the effects of DNA linker length, histone tail acetylation, linker histone H1, and multi-bromodomain proteins such as BRD4. It is designed to be experimentally accurate but computationally efficient, allowing the study of 100kb genomic regions on a timescale of seconds with moderate resources. We apply this model to explore the structure and dynamics of two active loci of mouse embryonic stem cells, *Pou5f1* and *Sox2*, as determined by solely their underlying epigenetic patterns. Our simulations reveal that chromatin folds into liquid-like domains characterized by similar histone modifications. These domains are highly dynamic, driving the formation of transient contacts between distant cis-regulatory regions. *In silico* mutation studies further clarify the distinct roles played by histone acetylation, linker histone, and BRD4. Overall, our physics-based modeling provides evidence that, in addition to other well-established mechanisms, epigenetic-dependent nucleosome-nucleosome interactions can play a key role in shaping the functional organization of genomic loci.

## INTRODUCTION

The hierarchical spatial organization of the Eukaryotic genome is critical for its function. Genomic DNA assembles with histone octamers to form nucleosomes, the basic structural unit of chromatin (∼10nm). Nucleosomes interact to form higher order structures including nucleosome clutches (∼20nm) (1,2), packing domains (PDs, 50∼200nm) (3,4), topologically associating domains (TADs) (5), and chromatin compartments (6). These domain structures are believed to aid the regulation of gene expression by controlling the assembly of the transcription machinery, bringing distant enhancers and promoters into proximity, or insulating genomic loci.

The Structural Maintenance of Chromosomes (SMC) complex cohesin has been recognized as a key player in the establishment of topologically associated domains through the mechanism of loop extrusion (7,8). However, the majority of TADs are maintained upon acute loss of loop extrusion factors (9), suggesting that additional mechanisms likely play a key role in chromatin 3D organization. The 1D pattern of epigenetic marks along the genome, such as histone tail acetylation, is another key factor long suspected to shape the 3D structure of the genome. The first Hi-C experiments already revealed how mammalian genomes are spatially organized into distinct large-scale compartments occupied by either active regions enriched in histone tail modifications (A compartment), or by inactive, gene-poor regions (B compartment) (6). Recent ultra-deep Hi-C data show that such organization persists down to the sub-TAD scale of individual genes, which are often organized into an alternating patterns of two chromatin compartments with an average interval size of 12.5 kilobases (10). The two compartments correlate with euchromatin/heterochromatin markers similarly to the original A/B compartmentalization, but on a smaller scale.

A recent experimental study measuring the condensability of native nucleosomes by next generation sequencing suggested that chromatin compartments are formed by phase separation through direct nucleosome-nucleosome interactions modulated by epigenetic modifications (11). This is consistent with the intrinsic tendency of chromatin to form complex epigenetic-dependent liquid condensates *in vitro* (12). These experiments suggest a regulation mechanism for *in vivo* chromatin domains, involving acetylation, linker histone H1, multi-bromodomain proteins and linker DNA length (controlling the spacing between nucleosomes along the genome). Despite this accumulating evidence, the extent to which epigenetics can shape the functional organization of active genomic loci in the absence of loop extrusion is still unclear.

Computer modelling provides suitable tools to address this question, as it allows to explore the dynamic organization of chromatin in controlled settings with high spatial and temporal resolution. In particular, coarse polymer simulations have been successfully employed to explore the roles of specific physical mechanisms shaping the 3D genome, such as SMC loop extrusion (8,13), bridging-induced attraction (14,15), epigenetic-driven phase separation (16–19), and nucleosome positions (20).

To unravel the role of epigenetics on the 3D organization of genes *in vivo*, we would ideally like to make predictions from the bottom up based on the physics of chromatin. There is a long history of all-atom and near-atomic coarse-grained molecular dynamics (MD) simulations that investigate nucleosomes (21–23), and nucleosome-nucleosome interactions (24,25). Coarser, but still physics based, nucleosome-level models have been successfully applied to explore the 3D structure and dynamics of chromatin (26–29), but sampling over sufficiently long timescales to observe the functional dynamics of genes remains a challenge.

In this work, we propose the Nucleosome Interaction Coarse Grained (NICG) model of chromatin, a physics-based model optimally designed to explore the epigenetic-dependent phase separation of chromatin into liquid domains. By representing proteins with one bead per histone and DNA with one bead per helix turn, our model has sufficient detail to capture the fundamental features of epigenetic-dependent nucleosome-nucleosome interactions, but efficient-enough to capture the dynamics of large (∼100kb) genomic loci over the timescale of seconds, allowing us to observe large-scale functional rearrangements of chromatin domains.

We applied our model to study the *Pou5f1* and *Sox2* loci of mouse embryonic stem cells (mESCs). We build a model of the target loci by incorporating key experimental epigenetic patterns that could potentially influence the phase separation of chromatin: nucleosome positions from chemical mapping (30), H3K27 acetylation ChIP-seq (31), histone H1 ChIP-seq (32), and multi-bromodomain protein BRD4 ChIP-seq (33). Our large-scale simulations reveal that the folding of the target loci is controlled by the underlying epigenetic profiles, with distinct roles played by acetylation, H1, and BRD4. These domains are highly dynamic and underly the formation of direct contacts between distant enhancers and promoters, showing direct evidence for a role of epigenetics in the functional organization of our genome.

## MATERIALS AND METHODS

### Nucleosome interaction coarse-grained model of chromatin

In our model, which we call the Nucleosome Interaction Coarse-Grained (NICG) model, we represent chromatin using solely point particles, with one bead for each histone protein in the nucleosome core, one bead for each B-form DNA helical turn (10.5bp) (34), and two beads for linker histone H1. Each nucleosome is constructed by arranging the 147 bp of nucleosomal DNA (=14 beads) along a superhelix with a radius of 40 Å and a pitch of 25 Å (35), and the 8 histone beads along a superhelix passing through the same axis and with the same pitch, but with a smaller radius of 18 Å (Figure 1A). Linker histone H1 is represented by two beads, one for the globular domain and one for the long, disordered C-terminal tail. When present, the H1 beads are positioned along the dyad, with the first bead 70 Å away from the nucleosome center. We employed a total of 20 bead types grouped into 5 classes characterized by the same interactions with others: linker DNA, canonical nucleosome, acetylated nucleosome, acetylated nucleosome associated with BRD4, and linker histone (Supplementary Table 1). All beads have the same size σ=35 Å, corresponding to the length of a DNA helical turn.

**Figure 1.**
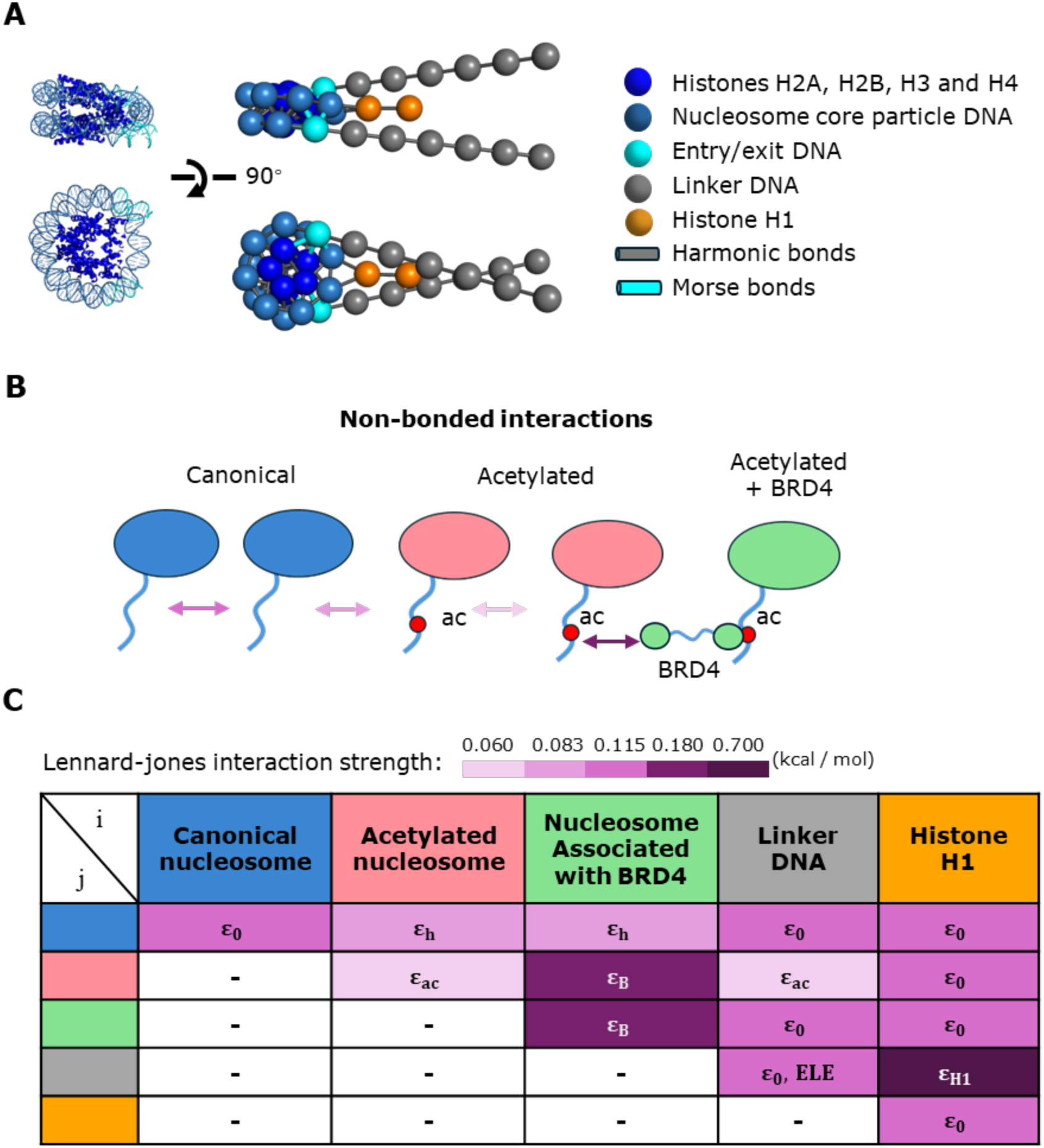
Architecture of coarse-grained chromatin model. **A.** Comparison between the X-ray crystallography structure (PDB ID :1KX5) and the coarse-grained model of a canonical nucleosome with linker DNA. In our model, we use one bead for each H2A, H2B, H3, or H4 histones, two beads for linker histone H1, and one bead for every DNA helical turn (10.5 bp). **B.** Schematics of non-bonded interactions between nucleosome beads and the mechanism of BRD4 bridging acetylated nucleosomes. **C.** Table of the non-bonded interaction strengths between the 5 groups of beads in our model. ε_0_, ε_h_, ε_ac_, ε_H1_: Lennard-Jones interaction strengths. ELE: Debye-Hückel electrostatic interaction. Relative strengths of Lennard-Jones interactions are indicated by color intensity.

The total potential energy is defined by contributions from local bond and angle potentials, and non-local Lennard-Jones and Debye-Hückel electrostatic interactions. Harmonic bonds are applied between all pairs of beads closer than 38 Å in the reference nucleosome geometry, and between consecutive linker DNA beads (Figure 1A). The first and last nucleosomal DNA beads (entry/exit DNA) was specially treated, and connected to the neighboring core beads by Morse bonds instead of harmonic bonds (Supplementary Information, Supplementary Figure S1A), which allows us to model the stochastic unwrapping of DNA from the histones (nucleosome breathing) (36,37). We applied angles potentials along the DNA chain to model its bending elasticity. Angle potentials are also applied between the first H1 bead and neighboring nucleosome beads to maintain the H1 globular domain dyad positioning (38). Our model does not account for DNA torsional elasticity and the effects of torsional stresses, reflecting the effects Type Ⅰ DNA topoisomerases (39), and also the accumulation of DNA twist within nucleosome core particles (40). Nevertheless, it is important to note that torsional stresses may be important in certain contexts (41), such as when studying the response of chromatin structure to gene transcription by RNA polymerase (42).

Non-bonded beads interact via a Lennar-Jones potential with strength varying according to their bead type, as described in Figure 1B-C. In particular, attraction was weakened for pairs of acetylated nucleosomes, but strengthened again for pairs where at least one acetylated nucleosome is associated with BRD4, to implicitly model the bridging effect of multi-bromodomain proteins (12,43) (Figure 1B). Interactions between linker DNA beads and H1 beads are strengthened to model H1-mediated compaction of nucleosome fibers. Linker DNA beads and H1 beads are also charged (-7.35e and +5e per bead, respectively) and they interact via an additional Debye-Hückel potential. Charge values are optimized to account for Manning condensation as well as possible within the considered framework of Debye-Hückel electrostatics.

### Coarse-grained simulation

All simulations were executed using the molecular dynamics software LAMMPS (44). Temperature was set to 300 K, while, except when noted, monovalent salt concentration was set to 150 mM to reflect physiological conditions. The equations of motion were integrated by Langevin dynamics. A uniform value of one unit mass was assigned to all bead types. Based on the analysis of chromatin diffusion (see section “Chromatin forms highly dynamic domains”), a scale factor of 1.4 ns/timestep was considered when comparing simulated time and real timescales. The Langevin damping factor was set at 140 ns, which corresponds to a low viscosity. Coordinates of all particles were saved every 10^5^ timesteps for further analysis.

### Force field parametrization and validation

The force field was parametrized based on a series of experimental data on chromatin organization: DNA persistence length, nucleosome unwrapping, and chromatin fiber sedimentation coefficients. It was then validated on experimental chromatin liquid-liquid phase separation properties.

First, we simulated a 1050 bp DNA molecule (100 beads) at various salt concentrations ranging from 2mM to 500mM, showing that we can reproduce the experimental DNA persistence length (Figure 2A) (45). These simulations were used to parametrize the DNA angle potential and electric charge.

**Figure 2.**
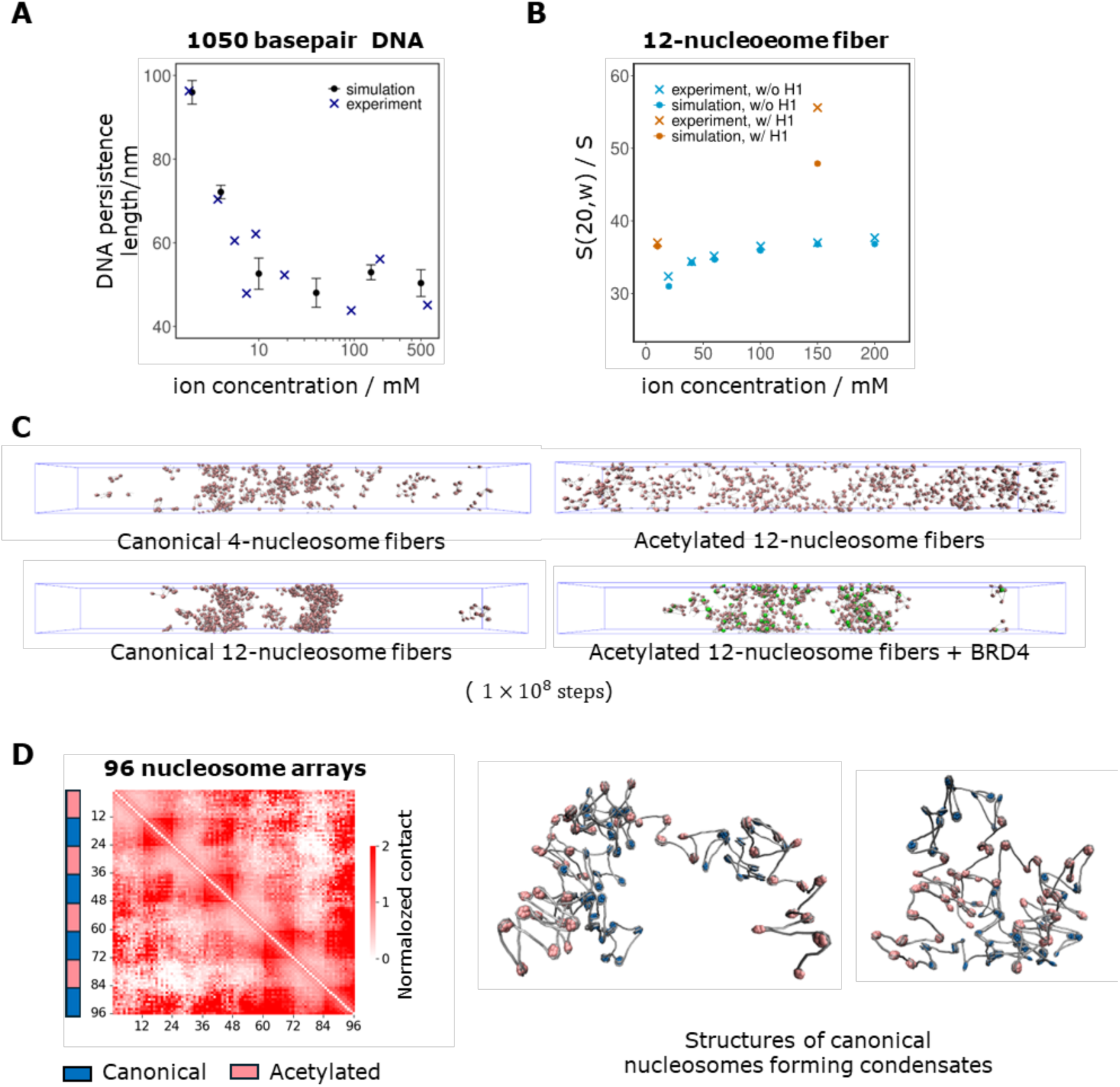
Model parameterization and validation. **A.** DNA persistence length as a function of salt concentration calculated from 1050 bp DNA simulations, compared with experimental data from Baumann et al.(45). **B.** Sedimentation coefficient of canonical 12-nucleosome fibers and canonical 12-nucleosome fibers with linker histone H1 as a function of salt concentration calculated from simulations, compared with experimental data from Hansen et al. and Grigoryev et al. (49,73). **C.** Slab simulation snapshots of 90 copies of canonical 4-nucleosome fibers or 45 copies of 12-nucleosome fibers with different modifications at the end of 10^8^ MD timestep. **D.** Normalized self-contact frequency map of 96-nucleosome arrays of alternating canonical and acetylated nucleosomes (left), and representative structure snapshots showing the condensation of canonical nucleosomes (right).

To optimize the Morse potential at the nucleosomal DNA entry/exit beads, we simulated 2 nucleosomes connected by a 63-bp linker DNA. The calculated equilibrium constant of spontaneous nucleosome unwrapping, 0.096 ± 0.006, was consistent with the experimental value at physiological conditions (0.02-0.1) (46). For acetylated nucleosomes, the unwrapping equilibrium constant from simulations is 0.32 ± 0.02, a ∼3 fold increase relative to canonical ones (Supplementary Figure S1B). Our model represents an “average” acetylated nucleosome with multiple acetylations on histone tails, so this is consistent with experiments showing that individual H3 or H4 acetylations increase the unwrapping probability by ∼2 fold (47).

Most of the remaining non-bonded interactions (the Lennard-Jones strengths ε, summarized in Figure 1C, and the H1 charge) are parametrized to reproduce salt-dependent sedimentation coefficients *s*_20,w_ of chromatin fibers (48,49). We performed simulations of 12-nucleosome fibers with 207-bp nucleosome repeat length (NRL) in the presence and absence of linker histone H1, in salt concentrations ranging from 10mM to 200mM (Figure 2B). The measured sedimentation coefficients are in good agreement with experimental data, except for some discrepancy at 150 mM salt in the presence of H1. This behavior is however consistent with the predictions from other coarse-grained models of chromatin (49), and we suspect that this may have to do with limitations in the prediction of the sedimentation coefficient from simulations in this regime (50). For acetylated nucleosomes, the Lennard-Jones interaction strength ε_ac_ was reduced to about half of ε_0_ (from 0.115 kcal/mol to 0.06 kcal/mol) to match experiments showing that acetylation reduces the sedimentation coefficients by a factor of 11-15% under physiological salt concentrations (51,52). This reduction in interaction strength is consistent with the change in nucleosome stacking free energy profiles measured experimentally by Funke et al. (53). The hybrid Lennard-Jones interaction strength between one bead from a canonical nucleosome core particle and one bead from an acetylated nucleosome core particle, ε_h_, is set according to the geometric average mixing rule. When present, linker histone beads interact with linker DNA beads via both electrostatic and Lennard-Jones interactions, the latter with enhanced strength ε_H1_. See Supplementary Information section on model parametrization for more details.

We validated our overall model through large-scale simulations of chromatin liquid-liquid phase separation using a slab geometric with a 100×100×1000nm^3^ simulation box (Figure 2C), under conditions where chromatin fibers are experimentally known to condense or not (12). As a negative control, we placed within the simulation box 90 copies of 4-nucleosome fibers; consistent with experiments, these short fibers did not phase separate in our simulations. We then placed 45 Copies of 12-nucleosome fibers in the same box, giving an initial nucleosome concentration of 90 μM. In this case the fibers phase separated into a dilute phase with a 12±1 μM and a condensed phase with a concentration of 294±12 μM, (Supplementary Figure S2A), which is consistent with the experimental observation of a condensed phase with a concentration of 342 μM. We do note some discrepancy with the experimental concentration of the dilute phase (32 nM), indicating that phase separation would occur earlier in experiments. In any case, both experiments and simulations show phase separations of unmodified 12-nucleosome fibers starting from concentrations close to that of the nucleus (∼100 μM), which is the regime we are most interested in. When we then placed 45 copies of acetylated 12-nucleosome fibers in the same box, we did not observe phase separation, consistent with experiments. Finally, we placed in the box 45 copies of acetylated 12-nucleosome fibers, where on each fiber 2 nucleosomes are also associated with BRD4. In this simulation, acetylated nucleosomes associated with BRD4 interact with other acetylated nucleosomes with enhanced Lennard-Jones strength ε_B_ (whether these are bound to BRD4 or not). In agreement with experiments, BRD4 association recovers the condensation of acetylated fibers (Figure 2C), confirmed by a higher number of nucleosome-nucleosome contacts (Supplementary Figure S2B).

As a final validation, we simulated a 96-nucleosome array alternating between canonical and acetylated every 12 nucleosomes (Figure 2D), as investigated in recent *in vitro* experiments (54). The resulting contact maps show a checkerboard-like pattern similar to those of experiments, with high contact frequencies between pairs of canonical nucleosomes, and lower contact frequencies between acetylated and other nucleosomes (Figure 2D, left panel).

### *Pou5f1* and *Sox2* loci model

The initial structures for the simulations of the *Pou5f1* and *Sox2* loci of mouse embryonic stem cells (mESCs) were prepared based on experimental information on nucleosome positions from chemical mapping (GSM2183909) (30), and epigenetic profiles from ChIP-Seq data (Figure 3, Supplementary Information methods). Acetylated nucleosomes are assigned according to the H3K27ac ChIP-seq profile in mESCs (GSM3399478) (31). Linker histone H1 is added to nucleosome dyad sites according to their H1 ChIP-seq profile (GSM1199586). BRD4 is assigned to acetylated nucleosomes according to the BRD4 ChIP-seq profile (GSM2319260) (33). See Supplementary Information section “Mapping of Epigenetic Patterns” for more details.

**Figure 3.**
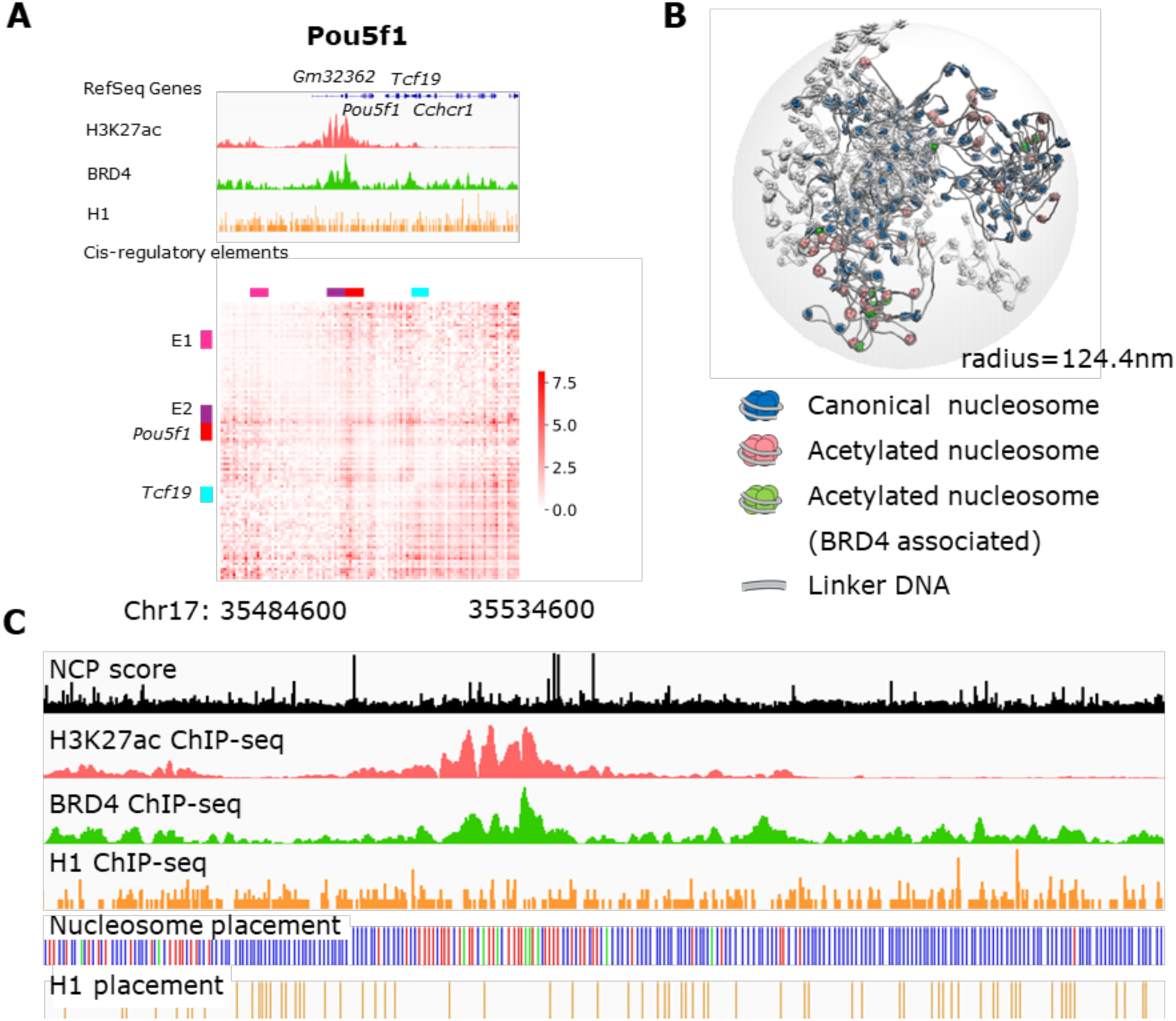
Coarse-grained modeling of the *Pou5f1* locus. **A.** Sequencing data of nucleosome interaction regulatory factors (H3K27ac, BRD4, and H1) and normalized Micro-C contact frequency map of the target genomic loci(58). **B.** Initial structures of the *Pou5f1* locus after a short relaxation by MD. White nucleosomes and DNA indicate the short chromatin fibers that are not the target locus. Canonical, acetylated, and BRD4-associated nucleosomes in blue, pink, and green, respectively. **C.** Mapping of nucleosome positions and modifications according to sequencing data (nucleosome colors as in panel B).

To recreate the chromatin density of mESCs nuclei, the spherical simulation space is filled with 5 short (10-16 kb) chromatin fibers in addition to the target locus, so that the average DNA concentration in the simulation space is 12.4 Mbp/μm^3^ (and the nucleosome concentration is ∼100 μM), as expected assuming the same DNA concentration as that in a mouse (5.4 Gbp) nucleus with a radius of 4.7 μm (55). Under the assumption that the chromatin environment surrounding these active euchromatin loci are euchromatin regions of similar composition as the considered loci, we assign 20% of the nucleosomes as acetylated, 31% of the nucleosomes as H1-bound, and 16.7% of the acetylated nucleosomes as BRD4-bound (the average values estimated from the number of canonical, acetylated, and BRD4-bound nucleosomes in the *Pou5f1* and *Sox2* loci).

The initial structures of the respective loci were generated by growing chromatin fibers that follows random reference backbones within spheres of radius 124.4 nm for *Pou5f1* and radius 156.7 nm for *Sox2* (more details in Supplementary Information and Supplementary Figure S4). Before the production runs, the initial structures, which may contain steric clashes, were first relaxed by molecular dynamics simulations using a soft-core potential and then with a short Langevin simulation using the full potential. Details on the contact map, structure clustering and domain analysis of the *Pou5f1* and *Sox2* simulations are provided in the SI.

## RESULTS

### Modeling of the mESC *Pou5f1* locus

To gain insight into the mechanisms that regulate chromatin structure *in vivo* at the sub-TAD scale, we applied our nucleosome-resolution computational model to the 50kbp *Pou5f1* gene locus of mESCs, located at chr17:35,484.6-35,534.6kbp (mm10) (56). In mESCs, the *Pou5f1* gene actively expresses a transcription factor key to the maintenance of pluripotency (57). This 50kbp locus alternates between high and low nucleosome acetylation (31), and the acetylated/non-acetylated intervals form sub-TAD self-interacting domains (10s of kilobases) on the experimental Micro-C contact map (58) (Figure 3A, Supplementary Figure S3). At this locus, contacts between distant acetylated enhancers and promoters are observed. Ensembl Regulatory features database (59) identified the *Pou5f1* promoter at chr17: 35506018 (P1). The locus also contains the active promoter of the *Tcf19* gene, which is colocalized with a small acetylation peak at chr17:35,516.8kbp (P2, +11kbp from P1). Groups of strong enhancers that contact the two promoters are located at chr17:35,485.6-35,486.1kbp (E1, -20kbp from P1) and chr17:35,502.1-35,502.7kbp (E2, -4 kbp from P1). Our simulations account for the modulation of nucleosome-nucleosome interactions by nucleosome positions, histone tail acetylation, linker histone H1, and BRD4 association. Our key focus is understanding to what extent epigenetic-dependent nucleosome interactions can play a role in the 3D organization of the considered locus, and if so, whether they can partially explain the observed contact maps. While cohesin is a key factor involved in the establishment of TADs, it is not explicitly modeled in our simulations as it is not the focus of our investigation, but we will comment on its potential role at the target loci.

The simulations were conducted in a spherical space with non-periodic boundaries, to mimic an isotropic confined nucleus environment. The space was supplemented with short, non-specific chromatin fibers to recreate the average DNA concentration in mESC cell nuclei. One initial structure after short relaxation simulations is shown in Figure 3B. Nucleosome positions, linker DNA lengths, distributions of canonical and acetylated nucleosomes, histone H1 and BRD4 were decided by incorporating experimental sequencing data (30,31,33) (Figure 3C).

### Nucleosome interactions induce 10-kilobase scale compartmentalization

We simulated 9 replicas of the *Pou5f1* locus using different initial structures, each for 0.7 seconds (5×10^8^ timesteps). The simulation successfully captured dynamical fluctuations of chromatin structures (Movie 1). The radii of gyration (Rg) in different replicas had a mean value of 74.8 nm, with a relatively low standard deviation of 5.2 nm (Figure 4A). The physical size of N-kbp continuous regions within the Pou5f1 locus scaled as N^1/D^, with D = 2.9 (Supplementary Figure S5A), consistent with experiments (60,61) and the theory of crumpled globules (62). We computed the contact frequency map from 9 simulation replicas using the structures from *t* = 0 to *t* = τ, and calculated correlation between independent replicas as a function of τ. The convergence of both R_g_ and contact map indicate simulations reached equilibrium (Supplementary Figure S5B). Even in the absence of cohesin, the simulated *Pou5f1* locus clearly separated into 3 domains consistent with acetylation profile (Figure 4B): the domain with intermediate level of acetylation (from -21 to -6 kbp), the domain with high level of acetylation (-5 to -3 kbp), and the domain with low level of acetylation (4 to 29 kbp). Each domain had high self-contact frequency. Principal component analysis (PCA) analysis of the normalized contact map identifies 2 compartments, characterized by either high or low acetylation levels, with PC1 being correlated with acetylation (Figure 4B, r = 0.55, p = 1.3×10^-10^). These results indicate that epigenetic modifications can play a significant role in the organization of the *Pou5f1* locus.

**Figure 4.**
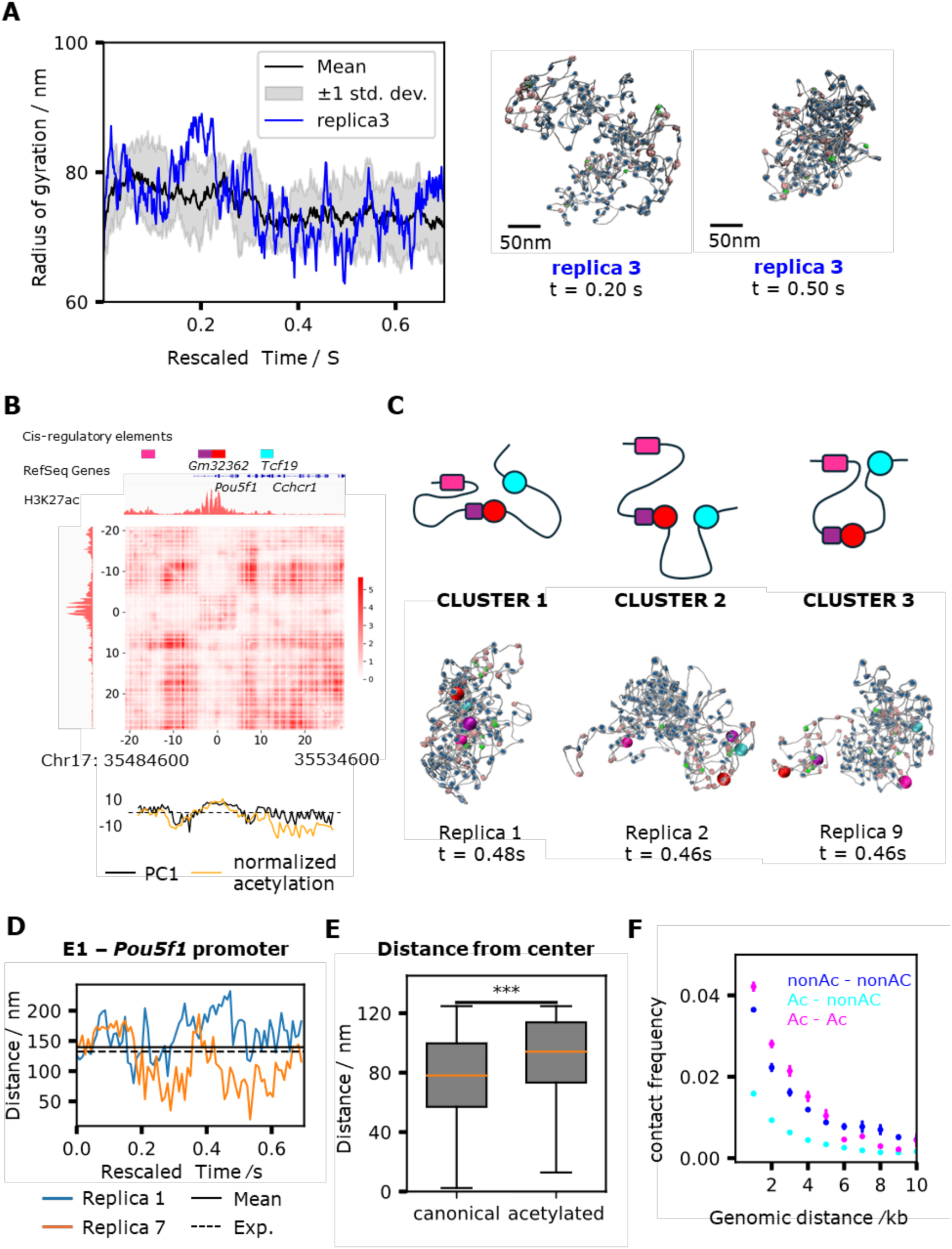
Nucleosome modification-mediated compartmentalization of *Pou5f1* locus. **A.** Mean radius of gyration (R_g_) over replicas and R_g_ in one representative replica of simulated models of the *Pou5f1* locus. Representative snapshots of open (Replica 3, t=0.20, R_g_=88.5 nm) and compact (Replica 3, t=0.50, R_g_=63.1 nm) structures are shown. **B.** Contact map calculated from *Pou5f1* locus simulation trajectories, projection of genomic position on the first component from principal component analysis of the contact map (PC1) compared with H3k27ac ChIP-seq signal. Data from t=0.14s to t=0.7s in 9 replica simulations are used. Acetylation was normalized by taking the logarithm of signal in each bin divided by mean signal of all bins, multiplied by a factor for better visual comparison. **C.** (Top) Illustration of regulatory element interactions in three major structure clusters. (Bottom) representative structure snapshots from each cluster. Regulatory elements E1 (-20kbp), E2 (-4kbp), P1 (0kbp), P2 (13kbp) are colored magenta, purple, red and cyan. **D.** Trajectories of distance between enhancer E1 and the *Pouf51* promoter (P1) in two representative replicas. Horizontal line indicates mean distances calculated from 9 replicas. **E.** Distribution of canonical and acetylated nucleosomes’ distances from the center of the spherical simulation box. **F.** Genomic distance dependent contact frequency between a pair of canonical-canonical, canonical-acetylated, or acetylated-acetylated nucleosomes in the *Pou5f1* locus. Data from t=0.14s to t=0.7s in 9 replica simulations are used. Standard error was calculated using data from 9 replicas as independent samples. Pairs with distance above 10kb were discarded because the sample size for acetylated pairs with genomic distance larger than 10kb was very small.

We also asked whether epigenetics alone could explain the experimental *Pou5f1* contact map from Micro-C data. PC1 calculated using normalized experimental Micro-C data showed visual correlation with simulation (Supplementary Figure S6), but the correlation is not statistically significant (r = 0.08, p = 0.38). In simulation contact map, the insulation scores at compartment boundaries had strong negative values, suggesting strong insulation between compartments, whereas the insulation scores calculated from experimental data were less profound (Supplementary Figure S6). The compartmentalization coefficient in simulation was 0.44, while 0.37 in experiment. This suggests that in the *Pou5f1* simulation, the phase separation of nucleosomes depending on acetylation, H1, and BRD4 was the primary force facilitating sub-TAD compartmentalization, while loop extrusion, transcription elongation and other factors acting *in vivo* but absent in our model could counteract this force. For example, in the experimental contact map a loop extrusion “stripe” brings the cohesin and CTCF binding site at 21.5kbp from the start of the locus in contact with the entire locus (Supplementary Figure S3A), blending the boundaries between 2 compartments, but this feature is absent in our simulations.

Direct communication between the two promoters and 2 enhancers mediated by nucleosome interactions were also observed in our simulations (Figure 4C). The distance between the first enhancer and the *Pou5f1* promoter (E1 and P1, genomic distance 20 kbp) had a mean value of 140 ± 5 nm across replicas, comparable with the experimental value of 132 nm (63) (Figure 4D). The standard deviation was 41 nm, while the experimental value was 84 nm. The additional heterogeneity *in vivo* may possibly be the result of heterogeneity at the level of epigenetic marks, which is absent in our model. K-means clustering of structures based on 6 pair-distances between regulatory elements (E1-E2, E1-P1, E1-P2, E2-P1, E2-P2, P1-P2) identified 3 major clusters of structures (Figure 4C, Supplementary Figure S7): Three major clusters were identified: in cluster 1, all regulatory elements interact in an overall compact chromatin structure; in clusters 2, E2, P1 and P2 interact, mediated by the clustering of acetylated nucleosomes; in clusters 3, E1 and P2 interact, mediated by the clustering of acetylated nucleosomes. The E1-P1 distances were largely different across replicas (Figure 4D), indicating the existence of different stable conformational states that each replica can adopt. Enhancer promoter contacts appear to be stabilized by the clustering of acetylated nucleosomes, in part associated with BRD4 (which mediates these interactions by bridging, representative snapshots shown in Figure 4C).

To gain insights into the overall organization of the *Pou5f1* locus, we calculated the distributions of distances of canonical and acetylated nucleosomes from the center of the spherical simulation space (Figure 4E, snapshots in Supplementary Figure S8). Interestingly, we find that canonical nucleosomes tend to occupy the core of the locus, while acetylated ones predominantly localize on the surface. This is in agreement with recent experiments and theoretical models (60,64) suggesting that heterochromatin-like compact chromatin locate at the core of domains, while euchromatin-like active chromatin locate at the periphery, where the transcription machinery is recruited.

Analysis of the contact frequency between nucleosome pairs as a function of their genomic distance confirms the role of epigenetics in compartmentalization: pairs of nucleosomes with the same chemical modifications (canonical-canonical or acetylated-acetylated) showed higher contact frequency compared to pairs with different modifications (Figure 4F).

### Chromatin forms highly dynamic domains

We next studied the dynamics of 10-kbp-scale domains involved in compartmentalization. For this analysis, we extended one *Pou5f1* simulation to 1×10^9^ timesteps (1.4 seconds). We find that despite forming well-defined compartments, single nucleosomes (both canonical and acetylated) diffuse over a vast space on our timescale of seconds, suggesting liquid-like, highly flexible chromatin structures (Figure 5A).

**Figure 5.**
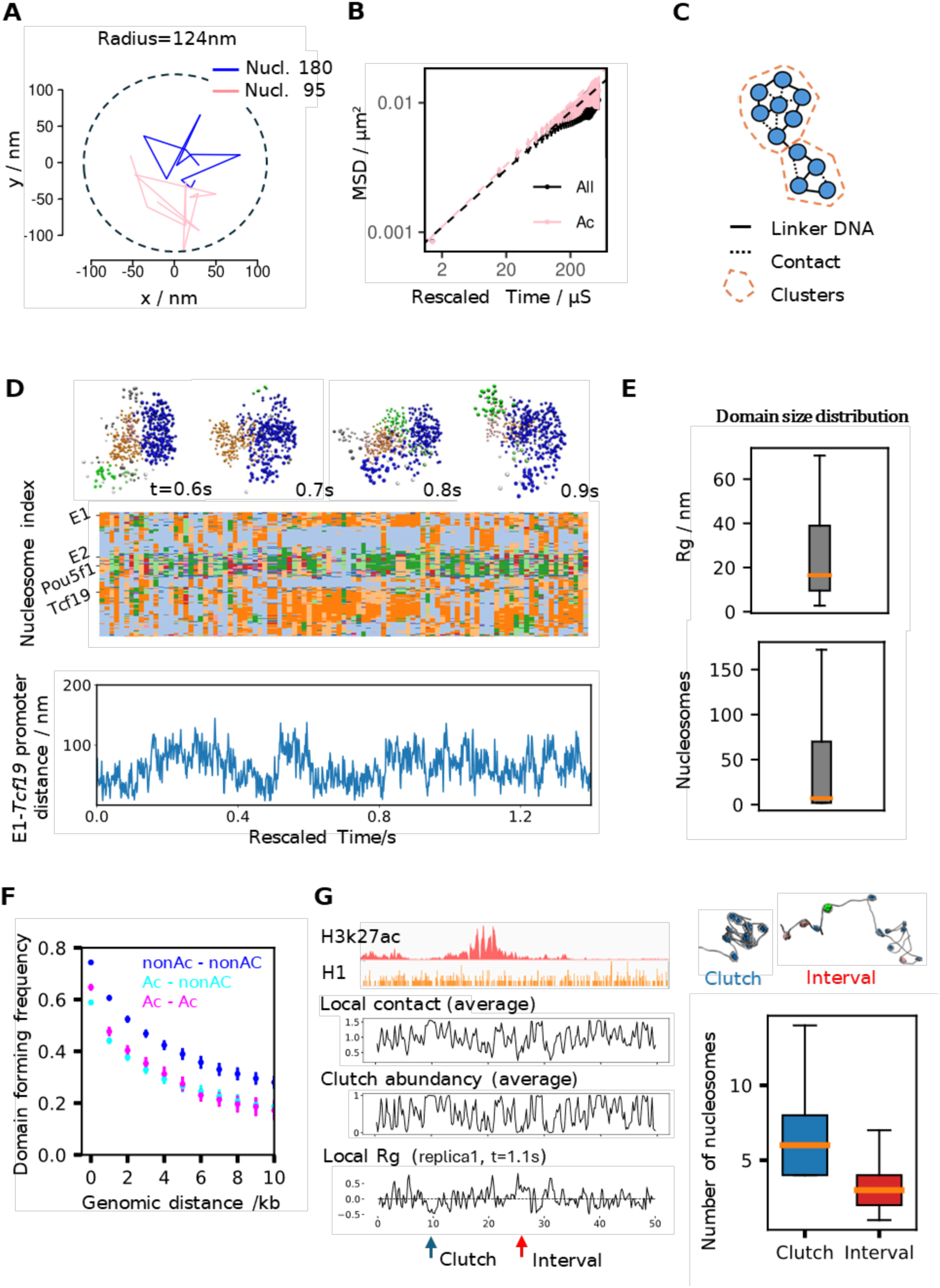
Multiscale dynamics of *Pou5f1* locus. **A.** Trajectories of the 180th nucleosome (canonical) and the 95th nucleosome (acetylated) from the extended simulation of the *Pou5f1* locus. Coordinates were taken every 0.14s. **B.** Mean squared displacement of all nucleosomes and acetylated nucleosomes in the *Pou5f1* locus, as a function of lag time. Dashed lines are fittings to linear model. **C.** Illustration of Infomap clustering applied to our chromatin system. Nucleosomes are the nodes in blue. **D.** (top) Representative snapshots of chromatin domains in the *Pou5f1* locus. Each nucleosome is visualized as one bead. “Background” non-specific chromatin fibers are included in the visualization. (Middle) Time series of domains plotted every 0.014 s from the extended *Pou5f1* locus simulation. Domains are indicated by different colors. “Background” nucleosomes are not included in the figure. (Bottom) Trajectory of the Enhancer1-*Tcf19* Promoter distance. **E.** Distribution of Infomap domain sizes in terms of radius of gyration (top) and number of nucleosomes (bottom). **F.** Frequency of a pair of canonical-canonical, canonical-acetylated, acetylated-acetylated nucleosomes being in the same domain as a function of genomic distance. Standard error was calculated using data from 9 replicas as independent samples. Data from t=0.14s to t=0.7s were used to avoid the effects of initial structures, **G.** (Left) Local contact frequency, clutch abundance (probability of each nucleosome being in a clutch) and 4-nucleosomal R_g_ from one structure snapshot plotted on the *Pou5f1* locus. (Right) Representative structures of one nucleosome clutch and one interval between clutches. Distribution of number of nucleosomes in clutches and intervals.

The mean square displacement (MSD) of nucleosomes follows a power law relation MSD = *K*_α_τ^α^ at small-enough lag time τ, before a plateau due to confinement (Figure 5B). α was 0.45 for the average over all nucleosomes, and 0.48 for acetylated nucleosomes, consistent with experiments (65) (α = 0.45 for all nucleosomes, α = 0.52 for nucleosomes in active early replication foci). Given the same exponent α, we could estimate the scale factor of 1.4 ns/timestep to map simulation times to real times by comparing the generalized diffusion coefficient *K*_α_ between simulations and experiments.

To study the *Pou5f1* locus dynamics, we used the Infomap community detection algorithm (66) (Supplementary Information) to assign all nucleosomes in the simulation to domains according to their spatial proximity (Figure 5C). This was done for every simulation frame to characterized how these domains change over time. Figure 5D shows that while domains tend to be stable for about 0.4 seconds, chromatin is also dynamic, with several domain splitting and merging events observed over the course of our 1.4 second trajectory, with such events being correlated with the fluctuation of enhancer-promoter distance (Figure 5D). Other, shorter simulation trajectories also show significant dynamic transformations of chromatin (Supplementary Figure S9).

The radius of gyration R_g_ of the domains is highly variable (Figure 5E), ranging from about 10 nm, on the order of individual nucleosomes and small nucleosome clutches (1,2), up to 60 nm, comparable to the size of packing domains observed in experiments (3,4).

We calculated the probability of a pair of nucleosomes being in the same domain as a function of their genomic distances (Figure 5F). As for contact frequencies, nucleosomes with the same modification have a higher chance to associate into a domain, again indicating that epigenetics drives compartmentalization.

### Epigenetic-dependent nucleosomes clutches along the chromatin fiber

We investigated the organization of groups of consecutive interacting nucleosomes (clutches), based on the mean contact frequency along 4-nucleosome sliding windows (Figure 5F). When averaged over the simulation trajectory, the mean contact frequencies anticorrelated with acetylation level, as expected from their weaker interactions. When considering individual simulation frames, the R_g_ along 4-nucleosome windows alternates between intervals of high and low values: we consider a continuous region in which R_g_ remains below average as one nucleosome clutch. Based on the analysis of clutches in the second half of the 1.4-second *Pou5f1* trajectory, clutch sizes vary between 4 and 14 nucleosomes (with a mean of 6.9), which is consistent with experimental estimates (2). Intervals between clutches vary between 1 and 19 nucleosomes, with a mean of 3.2. The probability of being part of a clutch in the trajectory was correlated with low acetylation and H1 binding.

### Chromatin interaction regulatory factors are essential for hierarchical organization

To assess the role of each individual nucleosome epigenetic factor for the 3D organization of the *Pou5f1* locus, we performed 1.4-second simulations mimicking various experimental conditions using the following 3 models (Figure 6): 1. Δnucleosome interaction: all non-bonded interaction strengths involving canonical nucleosomes are reduced to ε = 0.06 kcal/mol to mimic a widespread histone acetylation (although these nucleosomes will not experience enhanced interactions with BRD4-bound nucleosomes), 2. ΔBRD4: BRD4-associated nucleosomes are replaced by acetylated nucleosomes without BRD4. 3. ΔH1: linker histone H1 is removed from the target *Pou5f1* locus, but there is still H1 in the short chromatin fibers in the environment.

**Figure 6.**
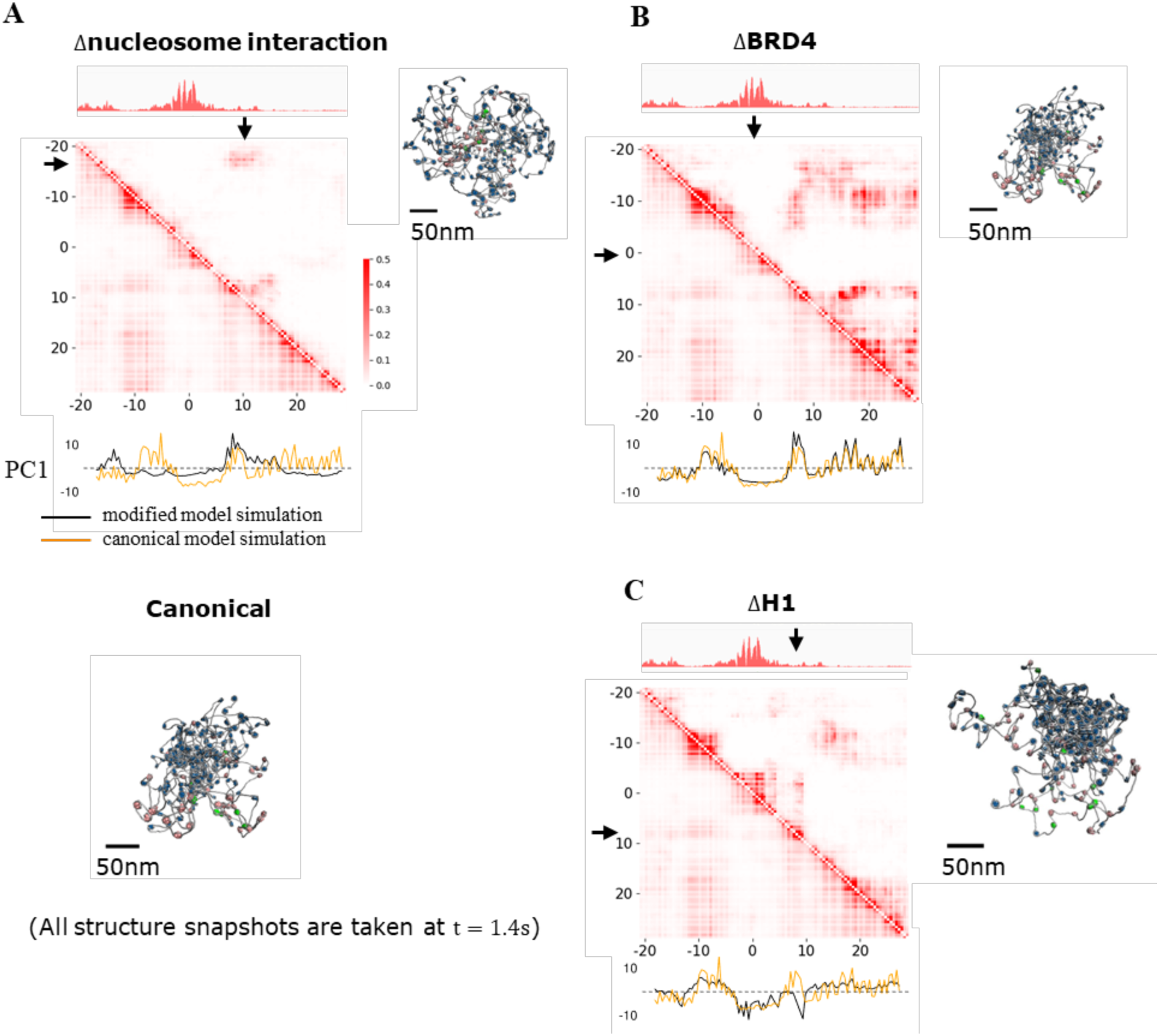
The impact of nucleosome interaction regulatory factors loss on chromatin compaction and compartmentalization. A∼C. Contact frequency maps and snapshots of the final structures of the modified *Pou5f1* locus with the final structure of the canonical *Pou5f1* locus as a reference. (A: interactions involving canonical nucleosomes are reduced. B: BRD4-associated nucleosomes are replaced by acetylated nucleosomes without BRD4. C: linker histone H1 is removed from the target *Pou5f1* locus). Below the contact maps we also show the projections of genomic loci on the first principal component from the canonical and the modified model simulations.

Reducing nucleosome interactions causes a reduction in the contact frequencies visible in the unnormalized contact map (Figure 6A), an increase in the physical size of the locus (Figure 7A), and significant changes in compartmentalization (Figure 6A). Due to the intact BRD4-mediated bridging between some of the acetylated nucleosomes, enriched interactions between enhancer E1 and *Pou5f1* promoter are still observed (arrows in Figure 6A), and the two acetylated sites are categorized as the same compartment by PC1.

**Figure 7.**
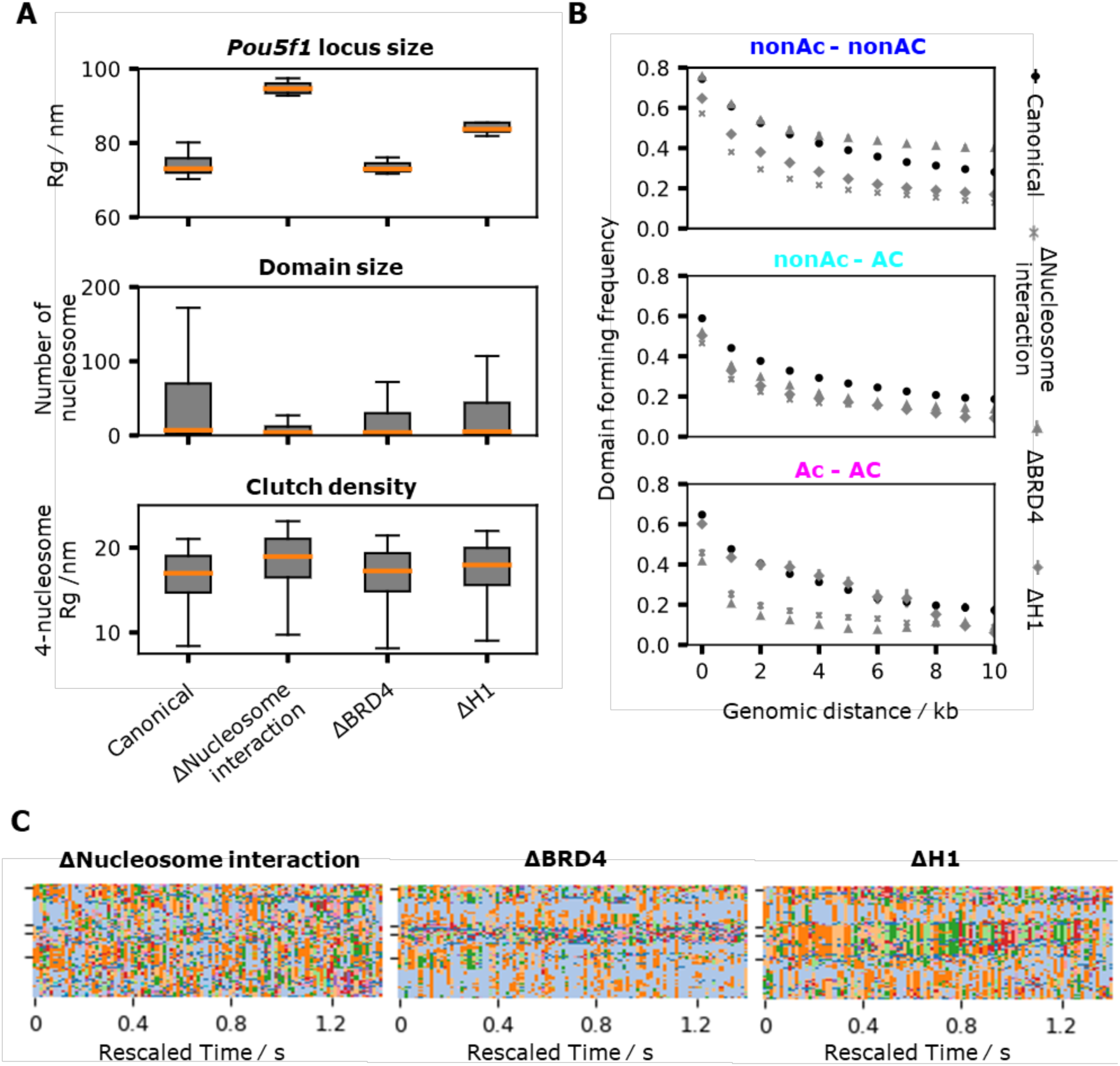
The impact of nucleosome interaction regulatory factors loss on chromatin compaction, domain formation and local chromatin fiber structures. **A.** (Top) Distribution of size (R_g_) in the canonical and modified *Pou5f1* locus simulations. (Middle) Distribution of Infomap domain size (number of nucleosomes) in the canonical and modified *Pou5f1* locus simulations. (Bottom) Distribution of R_g_ in regions identified as clutches in the canonical and modified *Pou5f1* locus simulations. **B.** Frequency of a pair of non-acetylated -non-acetylated (top), non-acetylated - acetylated (middle), acetylated-acetylated (bottom) nucleosomes in the canonical and modified *Pou5f1* locus simulations being in the same domain as a function of genomic distance. **C.** Time series of domains from the modified simulations indicated by different color. Plotted every 0.02s.

In the ΔBRD4 simulation, locus size (Figure 7A) and compartments (Figure 6B) remain largely unchanged compared to the unmodified model. However, the self-interacting domain at the -5 to 3 kbp acetylation peak is lost (Figure 6B), while E1-P1 and E2-P2 interactions are weaker. Repeating the same domain analysis as Figure 5D (Figure 7C), we found a decrease in association between acetylation peaks. This is also reflected in the probability of pairs of acetylated nucleosomes being in the same domain, which is lowest in the absence of BRD4 among the various conditions (Figure 7B). These results confirm the key role of BRD4 for the formation of active chromatin domains.

In the absence of H1, we observe an increase in *Pou5f1* locus size (Figure 7A) and a shift in domain formation near the moderately acetylated region at 4-9 kbp (Figure 6C). Instead of interacting with the upstream low-acetylation region between two acetylation peaks, this region forms contacts with its neighboring acetylation peak, shifting from the canonical compartment to the acetylated compartment (Figure 6C). This shift suggests a tug of war between different nucleosome interactions during compartmentalization: either mediated by H1 or by BRD4. The level of acetylation normally decides whether one genomic region will be in a mostly acetylated compartment or a mostly non-acetylated compartment, but H1 depletion disrupts the balance of the two mechanisms, resulting in compartment shifts. H1 depletion further causes a decrease in the mean number of nucleosomes within Infomap domains from ∼46 to ∼33 (Figure 7A). Clutches are also affected, with the mean value of 4-nucleosome R_g_ increasing to 17.7nm from 16.7nm (Figure 7A).

### Exploring the structure and dynamics of the 120-kbp *Sox2* locus

To test the capability of our physics-based nucleosome-resolution model on larger systems, we constructed the model of a 120 kbp region of mESCs, at chr3:34645kbp-34765kbp (mm10), containing the *Sox2* gene, simply referred to as the *Sox2* locus hereafter. *Sox2* is also a transcription factors and pluripotency regulator expressed in mESCs (57). Enriched contacts between two distal acetylated regions are observed in the experimental contact map (Supplementary Figure S3B). The *Sox2* promoter is located at chr3:34,650.4 kbp. Gene expression is controlled by a strong enhancer at chr3:34,756.5kbp-34,761.5kbp, ∼109 kbp downstream from the promoter (67). While enriched contacts colocalize with cohesin binding at this locus, we wanted to ask whether nucleosome-nucleosome interactions could partially contribute to enhancer-promoter communication at this locus in addition to loop extrusion. Using a setup analogous to that of *Pou5f1* (Figure 8A), we simulated 5 independent replicas of the *Sox2* locus for 0.42 seconds each (Movie 2).

**Figure 8.**
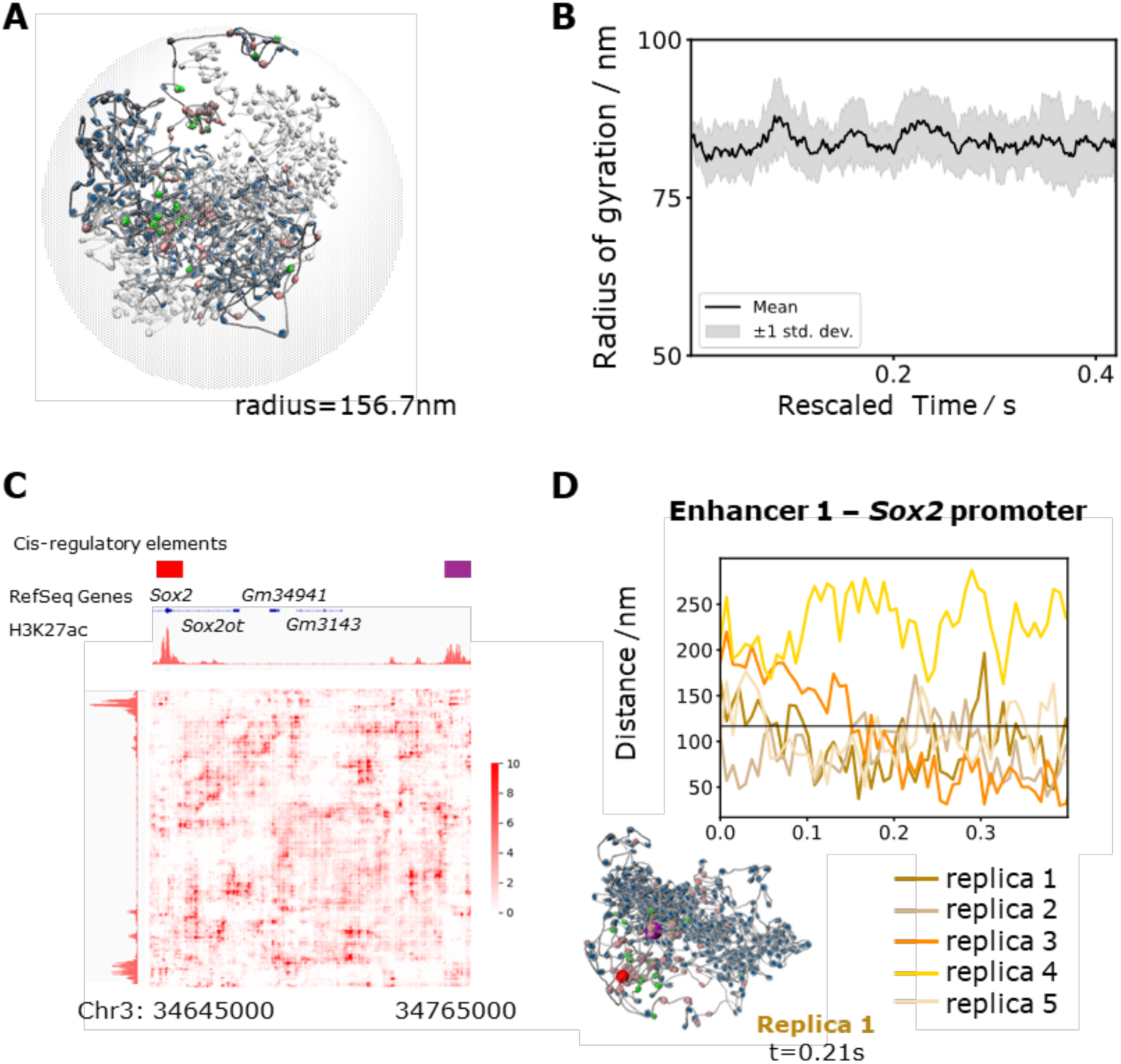
Simulation of the *Sox2* locus. **A.** Initial structure of the *Sox2* locus after a short relaxation simulation. **B.** Radius of gyration of 5 simulated replicas. **C.** Contact map calculated from t=0.2s to t=4.2s of 5 replicas of *Sox2* locus simulations. The Sox2 promoter and its strong enhancer are the acetylation peaks highlighted in red and purple, respectively. **D.** Trajectories of the enhancer-promoter distances in 5 replica simulations as a function of rescaled time, with one representative snapshot of enhancer-promoter interaction (promoter in red, enhancer in purple).

The mean R_g_ of the *Sox2* replicas was 83.8 nm, with a standard deviation of 4.7 nm (Figure 8B). Rg in each replica fluctuated around its mean value, showing convergence of this observable. Similar to the *Pou5f1* locus, the size of *Sox2* follows a power law with R_g_ ∝ N^1/3^ (Supplementary Figure S5B). Overall, the *Sox2* locus appears more compact compared to *Pou5f1*, perhaps due to the comparatively higher amount of H1 in this region.

There is no significant convergence between contact maps calculated from independent replicas (Supplementary Figure S5B), indicative of a lack of convergence. However, we do observe that in four out of five simulations the *Sox2* promoter form contacts with the 109 kbp downstream enhancer after 0.15 seconds (Figure 8D). Visual analysis of the structure suggests that these contacts are driven by interactions between acetylated nucleosomes mediated by BRD4.

## DISCUSSION

In this study, we designed and parameterized a nucleosome-resolution chromatin model, named NICG, capturing the effects of key epigenetic factors that modulate nucleosome-nucleosome interactions: linker histone H1, histone tail acetylation, and BRD4 association. We applied this model to explore the structural dynamics of the 50-kbp *Pou5f1* and 120-kbp *Sox2* loci of mESCs, focusing on the potential role of epigenetic-dependent nucleosome interactions to govern the 3D organization of these loci *in vivo*.

Our model is notable for being computationally efficient and physics-based, being rigorously parametrized based on a large set of experimental data including chromatin sedimentation coefficients and validated on chromatin liquid-liquid phase separation. The sole use of point particles and simple interaction potentials allows it to be easily implemented using most MD software (in our applications, LAMMPS (44)) and to make use of high-performance computing solutions. For example, our *Sox2* simulations included 200 kbp of chromatin and could run at a speed of 18 ms/day on a 112-core cluster. This allows us to study not only the structural organization of large genomic loci but also significant structural changes that occur over the timescale of seconds.

Our in-depth study of the *Pou5f1* locus of mESCs reveals that nucleosome-nucleosome interactions can play a significant role in the 3D organization of this gene. In particular, we find that the pattern of histone acetylation governs *Pou5f1* folding, with contact maps showing one heterochromatin-like compartments enriched in canonical nucleosomes often associated with linker histone H1, and another compartment including active enhancers that is enriched in acetylated nucleosomes sometimes associated with BRD4. Our *in silico* mutation studies clarify the distinct roles of separate epigenetic factors: acetylation weakens nucleosome interactions to expand chromatin, linker histone H1 is the main driver of compaction, while BRD4 bridges acetylated nucleosomes to bring together distant regulatory regions. We note that while these are not unexpected features given our knowledge of these factors, the simulations using our physics-based model suggest that epigenetics is likely to have strong effects on the genome organization *in vivo*. This agrees with a recent experimental study showing that the intrinsic strength nucleosome-nucleosome interactions as measured by their tendency to form condensates is sufficient to explain large-scale A/B compartmentalization (11). Our work expands on this to further show that these effects are likely relevant for sub-TAD compartmentalization as well (10). *In vivo*, epigenetic-driven nucleosome-nucleosome interactions likely act in concert with (or are antagonized by) other physical mechanisms such as cohesin loop extrusion (8). The role of nucleosome interactions could potentially explain puzzling experimental observations showing that contacts between promoters and distant enhancers are often minimally affected when cohesin activity is perturbed (9). According to our model, in the absence of loop extrusion enhancer-promoter communication could be rescued by interactions between acetylated nucleosomes bridged by BRD4, which can recover the phase separation of acetylated chromatin fibers *in vitro* (12).

Our simulations also offer insights into the functional organization of active loci such as *Pou5f1*. We find the typical organization of this locus consist of a heterochromatin-like core enriched in canonical nucleosomes surrounded by active acetylated regions that include enhancers and promoter, making these regions more accessible by the transcriptional machinery. This picture aligns with recent experimental and theoretical works (60,64): our work suggests that this organization naturally arises from intrinsic nucleosome interactions. The local structure of the chromatin fiber is also functionally organized into compact clutches typically enriched in canonical nucleosomes and H1, and separated by intervals of acetylated nucleosomes. The literature is rich in a variety of terms describing different levels of chromatin domains, but our analysis suggest that domains exist on a continuum, from small clutches (1) to intermediate packing domains (3) to larger compartments (6,10), and that nucleosome interactions alone are sufficient to give rise to such multi-scale organization.

Finally, our long simulations reveal the liquid-like nature of chromatin organization: despite domains being relatively stable, we observe several domain splitting and merging events over the timescale of seconds. Furthermore, we could also observe large changes in the distance between enhancers and promoter: which within 1 second may go from being 200-nm apart to come as close as 50 nm, which is sufficient to accommodate direct contacts mediated by the RNA polymerase-mediator complex. This suggest that enhancer-promoter communication is transient and occurs over timescales significantly faster than that of transcriptional bursting, which agrees with experiments and theoretical modelling (68). This also rationalizes the surprising observation that gene activation state appears to be mostly uncoupled from the underlying chromatin structure (69).

Our nucleosome-resolution model offers a flexible platform to explore the physical basis of the interplay between chromatin organization and gene function. While the current work focused on epigenetic-dependent nucleosome interactions, future work should aim to simulate the dynamics of a gene locus *in vivo* including additional physical processes that likely play a role to functionally organize the 3D genome (e.g., cohesin loop extrusion and transcription factor condensation) (70). Finally, our computationally lightweight model is ideally suited to be augmented with experimental biases (for instance from Micro-C data (58)) to implicitly model *in vivo* effects that cannot be directly captured by the physical model (71,72).

