## Supplementary Information for "Physics-based nucleosome-resolution modeling of epigenetic-driven chromatin domain dynamics"

**Details of model parametrization**

Particle types are listed and described in Supplementary Table 1. Force field form and parameters are listed in Supplementary Table 2.

1. DNA persistence length simulation

1050 base pairs of b-type DNA were modeled as a polymer of 100 beads, connected by harmonic bonds Lennard-Jones interaction and electrostatic interaction was applied between beads (supplementary table 1). Angel potential was applied to neighboring bonds. Simulations were run at salt concentrations 2mM, 4mM, 10mM, 40mM, 150mM and 500mM, each for $10^9$ steps.

DNA persistence length was calculated by calculating the autocorrelation (1) $C(n) = \langle \cos\ \theta_{i,i+n} \rangle$ of all bond vectors separated by n bonds. $C(n)$ is fitted to an exponential decay:

$$C(n) = \exp\ (l/l_p)$$

Where distance $l = nl_b$, $l_b$ is bond length, and $l_p$ is the persistence length. $l_p$ is calculated by fitting the $log\big(C(n)\big) \sim l$ relation to a linear model and taking the slope $b = l_p^{-1}$. The data for $l > 600$ nm is discarded to avoid finite length effect. The first $2 \times 10^8$ steps of the simulation under each condition are discarded to ensure simulation reaches equilibrium. The rest of the trajectory is divided into 4 parts, $l_p$ is calculated from each part. DNA persistence length is calculated as $\langle l_p \rangle$ and standard error was calculated as $\sigma(l_p)/\sqrt{4}$.

2. Nucleosome unwrapping simulation

Canonical nucleosome dimer was modeled as 2 canonical nucleosome core particles connected by 6 linker DNA beads (63 bp linker DNA). Equilibrium constant $K_{eq}$ was defined as the ratio of the probability of unwrapped states to the probability of the fully wrapped state (2).

Acetylated nucleosome dimer is modeled as the same geometry, with Lennard-Jones interaction strength between nucleosome core particle beads and Morse bond strength between the entry/exit DNA bead and nucleosome core particle beads set to the acetylated values (Supplementary Table 1).

Nucleosome dimers are simulated $1 \times 10^8$ steps at 150mM salt concentration. We calculated the distance $d$ between the exit DNA bead on the first nucleosome and the NCP-DNA bead 8 beads (approximately one turn of DNA wrapped on nucleosome) upstream, and defined structures where $d \leq d_0$ as wrapped states, structures where $d > d_0$ as unwrapped states. $d_0$ was set at 35nm. Each simulation trajectory was divided into 5 intervals, each $2 \times 10^7$ steps. $K_{eq}$ was calculated in the last 4 intervals to ensure simulation reached equilibrium. Standard error was calculated as $\sigma(K_{eq})/\sqrt{4}$.

3. Nucleosome fiber sedimentation coefficients simulation

Sedimentation coefficient standardized to conditions corresponding to pure water at 20 °C and extrapolated to zero protein concentration, $s_{20,w}$, was used to quantify the compaction of chromatin fibers (3–6).

Canonical 12-nucleosome fiber was modeled as 12 canonical nucleosome core particles connected by 11 linkers with lengths 6, 6, 5, 6, 6, 5, 6, 6, 5, 6, 6 beads. The linker DNA length in the considered experiments is 207-147=60bp ~ 5.7 coarse-grained DNA beads (5,7). There was no overhanging linker DNA on the outer side of the end nucleosomes. A single fiber was simulated at 20mM, 40mM, 60mM, 100mM, 150mM, 200mM salt concentrations, for $2 \times 10^8$ steps under each condition.

Acetylated 12-nucleosome fiber was modeled with the same geometry as canonical 12-nucleosome fiber, with Lennard-Jones interaction strengths and Morse bond strengths set to the acetylated values (Supplementary Table 1). A single fiber is simulated at salt concentration 150mM for $2 \times 10^8$ steps.

Canonical 12-nucleosome fiber with linker histone H1 was modeled as the canonical 12-nucleosome array with H1 placed at the dyad site of every nucleosome. A single fiber was simulated at salt concentrations 10mM and 150mM, for $2 \times 10^8$ steps under each condition.

$s_{20,w}$ wass calculated using the method described by G. Arya *et. al.* (8):

$$s_{20,w} = s_1 \left( 1 + \frac{2R}{N} \sum_{i}^{N} \sum_{j>i}^{N} \frac{1}{R_{ij}} \right)$$

Using the same value of $s_1$ ($s_{20,w}$ of one single nucleosome) and $R$ (diameter of one nucleosome) as in previous computational studies (3). $N$ is the number of nucleosomes and $R_{ij}$ is the distance between two nucleosomes.

$s_{20,w}$ was calculated for each structure snapshot. The first $1 \times 10^8$ steps of the simulation under each condition were discarded to ensure simulation reaches equilibrium. Autoregressive integrated moving average (ARIMA) analysis is performed to the time series of $s_{20,w}$ to determine the mean value and its error. ARIMA analysis was done using the R package "forecast"(9,10).

4. Nucleosome fibers phase separation simulations

4-nucleosome fibers were constructed as 4 nucleosomes connected by 4 beads linkers (42 bp linker length). 90 copies of the same fiber were placed in a $100\text{nm} \times 100\text{nm} \times 1000$ nm periodic boundary simulation box. Canonical 12-nucleosome fibers were constructed as 12 nucleosomes connected by 4 bead linkers. 45 copies of the same fiber were placed in a $100\text{nm} \times 100\text{nm} \times 1000$ nm periodic boundary simulation box. Acetylated 12-nucleosome fibers were constructed as 12 acetylated nucleosomes connected by 4 bead linkers. 45 copies of the same fiber were placed in a $100\text{nm} \times 100\text{nm} \times 1000$ nm periodic boundary simulation box. Acetylated 12-nucleosome fibers associated with BRD4 were modeled as 10 acetylated nucleosomes (not associated with BRD4) and 2 acetylated nucleosomes associated with BRD4, connected by 4 bead linkers. The BRD4-associated nucleosomes were placed at the second and the 11th nucleosomes. 45 copies of the same fiber were placed in a $100\text{nm} \times 100\text{nm} \times 1000$ nm periodic boundary simulation box.

Simulation for each type of nucleosome fiber was run for $1 \times 10^8$ steps, simulation convergence was confirmed by calculating the total number of nucleosome contacts (figure S2. Contact was counted when the distance between two nucleosome centers was smaller than 110 nm). The first $2 \times 10^7$ steps of each trajectory were discarded to ensure simulation reached equilibrium.

The one-dimensional nucleosome concentration was calculated along the long axis of the simulation box, the regions with a concentration higher than $0.05\ \text{nucleosome} / \text{Å}$ were assigned as the condense phase (Figure S2). The three-dimensional nucleosome concentration was calculated for each frame to calculate the mean concentration in the condense phase.

We used a BDR4 / acetylated nucleosome ratio (~0.17) close to the value in active gene loci. We obtain the ratio from the observed number of BRD4 molecules (9~15 molecules) at the Pou5f1 gene locus in mouse ES cells (mESCs) (11) and the estimated number of acetylated nucleosomes at the same locus (see section "Mapping of nucleosome position and nucleosome modifications ").

5. 96-nucleosome array simulation

A 96-nucleosome array was constructed as 96 nucleosomes connected by 95 linkers of 6 beads (63 bp linker length). The $(i \times 12 + 1)$ -th to the $((i + 1) \times 12)$ -th nucleosomes were acetylated nucleosomes, $i = 0, 2, 4, 6$ . The $(j \times 12 + 1)$ -th to the $((j + 1) \times 12)$ -th nucleosomes were canonical nucleosomes, $j = 1, 3, 5, 7$. 10 replicas of the same array were simulated for $1 \times 10^8$ steps with different random seeds. The first half of all trajectories were discarded to ensure simulation reached equilibrium. Contact frequency was calculated by normalizing contacts counted in all trajectories to the total amount of structure frames. Contact was defined by the sigmoid function described in the section "Contact map data analysis" in Supplementary methods.

**Mapping of nucleosome positions, nucleosome acetylation, linker histone H1 and multi-bromodomain protein BRD4**

The center of nucleosome core particles (nucleosome dyads) were mapped to the genome using the chemical mapping data of nucleosomes (12) (GEO accession number: GSM2183909). Genomic coordinates in each target locus (Pou5f1 and Sox2) were sorted by their chemical mapping score. We assigned nucleosomes to the coordinate with the highest score that is at least 146 bp from other already assigned nucleosomes and repeat this process until no more nucleosomes can be assigned. Linker length was decided by subtracting 146 bp from the distance between two neighboring nucleosome dyads. In the case where linker DNA length couldn't be represented by an integer number of beads, rounding up was applied to ensure at least one linker bead presented between two nucleosome core particles.

Acetylated nucleosomes were assigned to nucleosomes according to their H3K27ac ChIP-seq signals in mESCs (13) (GSM3399478). The ChIP-seq signal $S$ of one nucleosome was defined as the ChIP-seq signal integrated in the 146 bp interval centered at the nucleosome center position. The acetylation state of a nucleosome was decided statistically using a random number generator.

If the random number $q \leq S/S_0$, the nucleosome was assigned as acetylated. The normalization factor $S_0$ was calibrated by randomly sampling 10000 nucleosomes from the entire genome using the unique peaks of nucleosome chemical mapping score from GSM2183909. $S$ of sampled nucleosomes were calculated and $S_0$ was set at 100.0 so that 1% of sampled nucleosomes were assigned as acetylated, comparable with the values ranging from 1.2% to 7 % reported in different cell lines (14,15).

Linker histone H1 was added to nucleosome dyad sites according to their H1 ChIP-seq signals in mESCs (GSM1199586). The same approach to assigning acetylated nucleosomes is applied here, with the normalization factor set so that histone H1/nucleosome ratio in randomly sampled nucleosomes is 0.36, consistent with the values ranging from 0.36 to 0.46 reported in mESCs (16,17).

BRD4 associated nucleosomes were assigned according to BRD4 ChIP-seq signals (18) (GSM2319260). BRD4 was first mapped to nucleosomes using the same approach described above. From the mapped nucleosomes, only nucleosomes already assigned as acetylated nucleosomes were assigned as acetylated nucleosomes associated with BRD4 in the model of target loci. The normalization factor was set so that the number of BRD4 molecules was 10 in the Pou5f1 gene locus, consistent with the finding that the number of BRD4 molecules at the Pou5f1 transcription foci are 9~15 (11).

**Construction of Pou5f1 locus and Sox2 locus initial structures**

The sizes of the confined simulation spaces were decided so that the total amount of chromatin in the two setups were 100 kbp for Pou5f1and 200 kbp for Sox2. Radius of the simulation spaces were set at 124.4 nm for the Pou5f1 locus and 156.7 nm for the Sox2 locus.

To setup the initial structures, replicas of even more coarse-grained backbone structures were first simulated using LAMMPS. Chains of beads were simulated in the spherical confined space. Segments between two neighboring beads represented 1kb chromatin. Beads were connected by harmonic bonds with no angle potential, and a Lennard-Jones potential with cutoff (only repulsive interactions) was applied between beads. The bead sizes (bond length and repulsive potential cut off) were set at 22.5 nm. Chains representing the Pou5f1 locus (50 kbp) and 4 short chromatin fibers (each 10 kb) were placed in the 124.4 nm sphere. Chains representing the Sox2 locus (120 kbp) and 4 short chromatin fibers (each 16 kbp) were placed in the156.7 nm sphere.

Nucleosome resolution models of target loci were constructed by placing nucleosomes one by one (Figure S4). The nucleosome core particle model in Figure 1A was copied, rotated and translated so that the entry DNA bead was placed at the start of the backbone (for the first nucleosome in the chain) or at the end of the linker of the previous nucleosome. The axis of nucleosome was rotated to be parallel to the backbone. Linker DNA at the downstream side of the nucleosome was generated by placing beads representing linker DNA on the line extended from the last NCP-DNA bead to the exit DNA bead, with 35Å distances between beads. The number of linker DNA beads was decided using the linker DNA length decided in the previous section.

The nucleosome resolution chromatin structure then goes through a soft-core potential simulation using LAMMPS. The Lennard-Jones interaction strength was set at 0 at the beginning of the simulation and linearly increased over the course of 100000 timesteps until it reached the target values in Supplementary Table 1. Harmonic bond and angel potentials were set as in Supplementary Table 1 and electrostatic interaction were not applied in this simulation. This simulation allowed automatic adjustment of the initial structure to avoid clashes between nucleosomes. The structure was further relaxed using a Langevin simulation with the full model (parameters as in Supplementary Table 1) for $1 \times 10^7$ steps to produce the initial structures presented in Figure 3B.

**Contact map data analysis**

"Hi-C" like contact maps were generated by calculating the contact frequency between nucleosomes. Contact was determined by a sigmoid function:

$$c = \frac{1}{1 + \gamma^6}$$

Where $\gamma = (\frac{r - d_0}{r_0})$, $r$ is the distance between the center of two nucleosomes, $d_0$ is the nucleosome diameter, 110 Å and $r_0$ is set as 55 Å. Nucleosomes are mapped into 500 base pair bins. Contact frequency in each bin is normalized by distance-dependent average.

Compartments were called by calculating the principal component analysis (PCA) of normalized contact map. Compartmentalization coefficient is calculated as $\frac{C_{in} - C_{out}}{C_{in} + C_{out}}$, where $C_{in}$ is the sum of contact frequencies between bins in the same compartment, and $C_{out}$ is the sum of contact frequencies between bins in different compartments.

Insulation score of one bin was calculated as the sum of contact frequencies between upstream bins and downstream bins in a 20-bin window (10 kbp) centered at the bin of interest, normalized by the mean value of all windows.

**Visualization of experimental Micro-C data**

Micro-C data (19) (GSE130275) were exported using Juicebox (20). Bin size was set as 500 base pair and contact frequency in each bin was normalized using the observed/expected normalization option.

***Pou5f1* Structure clustering using K-means.**

Distances E1-E2, E1-P1, E1-P2, E2-P1, E2-P2, P1-P2 were calculated every frame from the t > 0.14s parts of all 9 replica simulations. K-means clustering was performed using the Python module pyEMMA (21) using different number of clusters k. Within-cluster-sum-of-squares (WCSS) was calculated for different k, and k=3 was chosen as the optimal number of clusters using the elbow criterion (Figure S7).

**Domain identification using Infomap Clustering**

Chromatin structures were converted into a network, with nucleosomes as nodes and nucleosome-nucleosome contacts as links. Contacts were identified using the same sigmoid function described in the section “Contact map data analysis” in Supplementary methods.

Clusters were optimized by minimizing the map equation (22). The map equation describes the conciseness of describing the trajectory of a random walk on the network by “movements between clusters” and “movements within clusters”. A python implementation of the algorithm was used (23). The depth level parameter was set as 1, meaning no sub-clusters were calculated. The sizes of clusters were automatically optimized.

After cluster calling for each structure frame, names were assigned to clusters frame by frame. To keep cluster names consistent throughout the time series, the name of cluster $A_i$ in frame $i$ will be inherited by $A_{i+1}$, the cluster with the most common members with cluster $A_i$, in frame $i+1$.

### Supplementary Tables

| Bead type | Name | Description |
|---|---|---|
| 1 | DLN | 10.5 bp linker DNA |
| 2 | DNC | 10.5 bp nucleosome core particle DNA not at the entry/exit site |
| 3 | DEX | 10.5 bp nucleosome core particle DNA at the entry/exit site |
| 4 | H2A | Histone H2A in a canonical nucleosome |
| 5 | H2B | Histone H2B in a canonical nucleosome |
| 6 | H3 | Histone H3 in a canonical nucleosome |
| 7 | H4 | Histone H4 in a canonical nucleosome |
| 8 | DAN | 10.5 bp nucleosome core particle DNA in an acetylated nucleosome, not at the entry/exit site |
| 9 | DAE | 10.5 bp nucleosome core particle DNA in an acetylated nucleosome, at the entry/exit site |
| 10 | A2A | Histone H2A in an acetylated nucleosome |
| 11 | A2B | Histone H2B in an acetylated nucleosome |
| 12 | A3 | Histone H3 in an acetylated nucleosome |
| 13 | A4 | Histone H4 in an acetylated nucleosome |
| 14 | DBN | 10.5 bp nucleosome core particle DNA in an acetylated nucleosome associated to a BRD4 molecule, not at the entry/exit site |
| 15 | DBE | 10.5 bp nucleosome core particle DNA in an acetylated nucleosome associated to a BRD4 molecule, at the entry/exit site |
| 16 | B2A | Histone H2A in an acetylated nucleosome bound to a BRD4 molecule |
| 17 | B2B | Histone H2B in an acetylated nucleosome bound to a BRD4 molecule |
| 18 | B3 | Histone H3 in an acetylated nucleosome bound to a BRD4 molecule |
| 19 | B4 | Histone H4 in an acetylated nucleosome bound to a BRD4 molecule |
| 20 | H1 | Linker Histone H1 |

**Supplementary Table 1.** Names and descriptions of bead types used in the nucleosome resolution chromatin model. Colors correspond to non-bonded interaction groups consistent with Figure 1 B-C in main text.

| Harmonic bonds $E = k(r - r_0)^2$ | | |
|---|---|---|
| $k = 0.50$ | | $r_0$= native value |
| **Morse bonds** $E = \mathrm{D}\left[1 - e^{-\alpha(r-r_0)}\right]^2$ | | |
| $\mathrm{D}_0 = 2.20$ <br> $\mathrm{D}_{ac} = 1.80$ | $\alpha = 0.3$ | $r_0$= native value |
| **Angle potentials** $E = K[1 + \cos(\theta)]$ | | |
| $K = 8.8$ | | |
| **Lennard-Jones interactions** $E = 4\varepsilon\left[\left(\frac{r}{\sigma}\right)^{12} - \left(\frac{r}{\sigma}\right)^{6}\right], r < r_c$ | | |
| $\varepsilon_0 = 0.115$ <br> $\varepsilon_{ac} = 0.060$ <br> $\varepsilon_h = 0.083$ <br> $\varepsilon_\mathrm{B} = 0.180$ <br> $\varepsilon_{H1} = 0.700$ | $\sigma = 35.0$ | $r_c = 79.538$ |
| **Debye-Hückel interactions** $E = \mathrm{C}\frac{\mathrm{q_i q_j}}{\epsilon r}\exp(-\mathrm{k}r), r < r_c$ | | |
| $\epsilon = 80$ | $k = 0.325\sqrt{\mathrm{I}}$ | $r_c = 10/k$ |

**Supplementary Table 2.** Pair interaction parameters used in the coarse-grained chromatin model. Distance units are Å, Energy units are $\mathrm{kcal \cdot mol^{-1}}$. $r_0$ for each pair of beads in the nucleosome core particle are set as in our reference model (see main text). $r_0$ for each pair of linker DNA beads are set at 35.0Å. For Debye-Hückel interactions, $k = \sqrt{\frac{\mathrm{I}}{\epsilon \mathrm{k_B T}}}$ is the inverse of Debye length (unit: $\mathrm{Å^{-1}}$). I is the ionic strength (unit: mol/L).

### Supplementary Figures

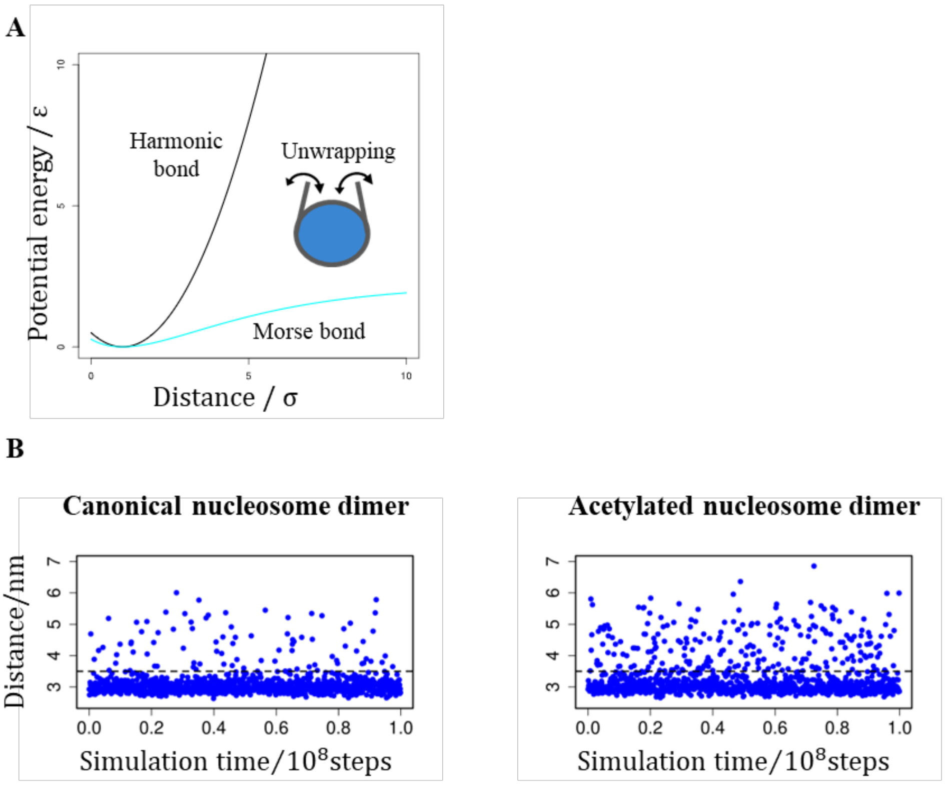


**Figure S1. A.** Comparison between the energy landscape of harmonic bonds and morse bonds. **B.** Simulation trajectory of the distance between the exit DNA bead and nucleosome core particle in a nucleosome dimer. Dashed line indicates the cutoff for unwrapped state, 3.5nm.

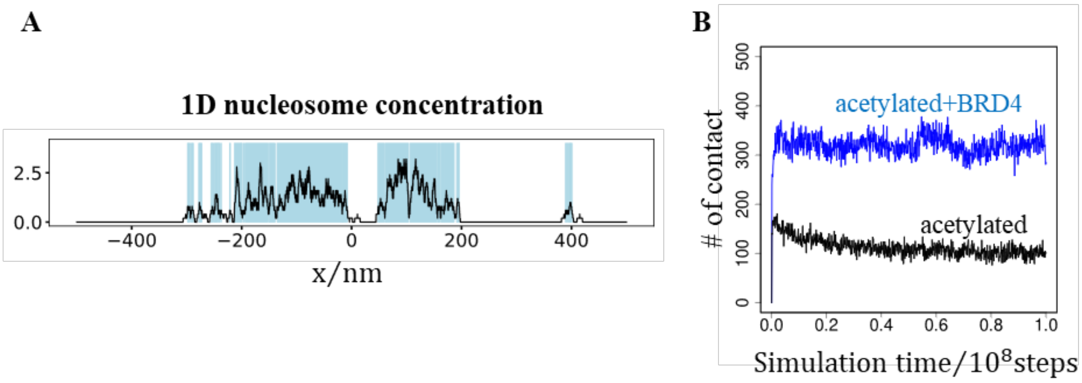


**Figure S2. A.** One dimensional nucleosome concentration at t=$10^8$ timesteps in the simulation of acetylated and BRD4 associated 12-nucleosome fibers. The unit of concentration is $Å^{-1}$. Regions highlighted in cyan are classified as condensates. **B.** Sum of nucleosome contacts in simulations of 12-nucleosome fibers.

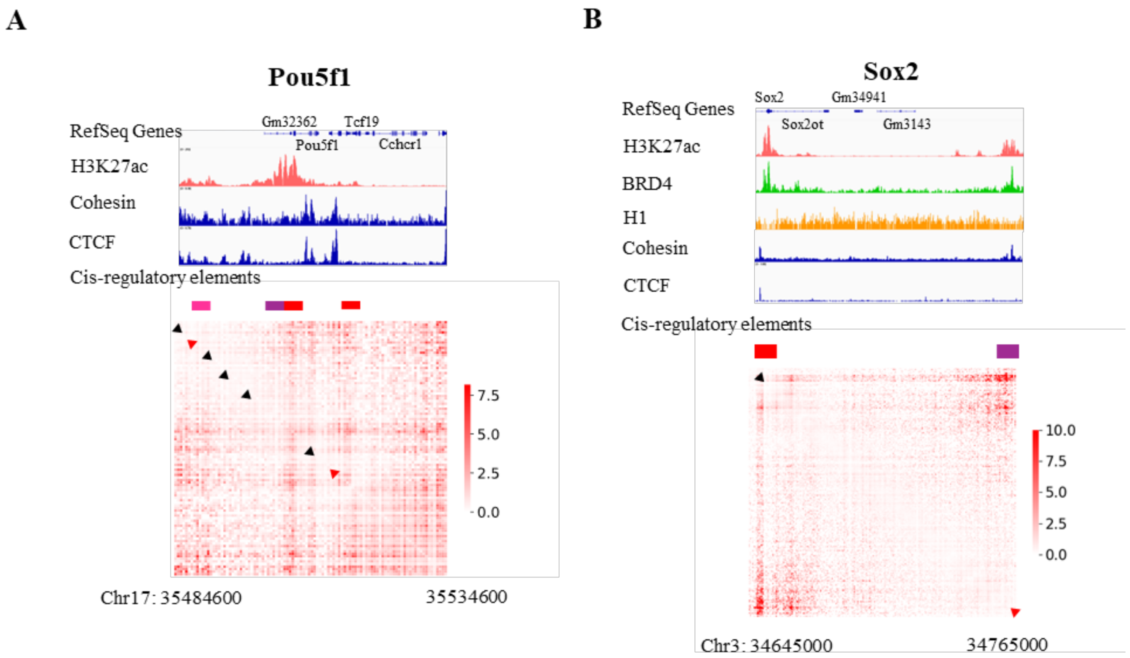


**Figure S3.** Normalized micro-C contact frequency data compared with ChIP-seq data. On the contact map, we also show the CTCF binding motifs scanned using FIMO (24). Motifs located on the positive strand and negative strand are colored black and red, respectively, with arrows indicating the inside of loops if loop extrusion stops at the corresponding site.

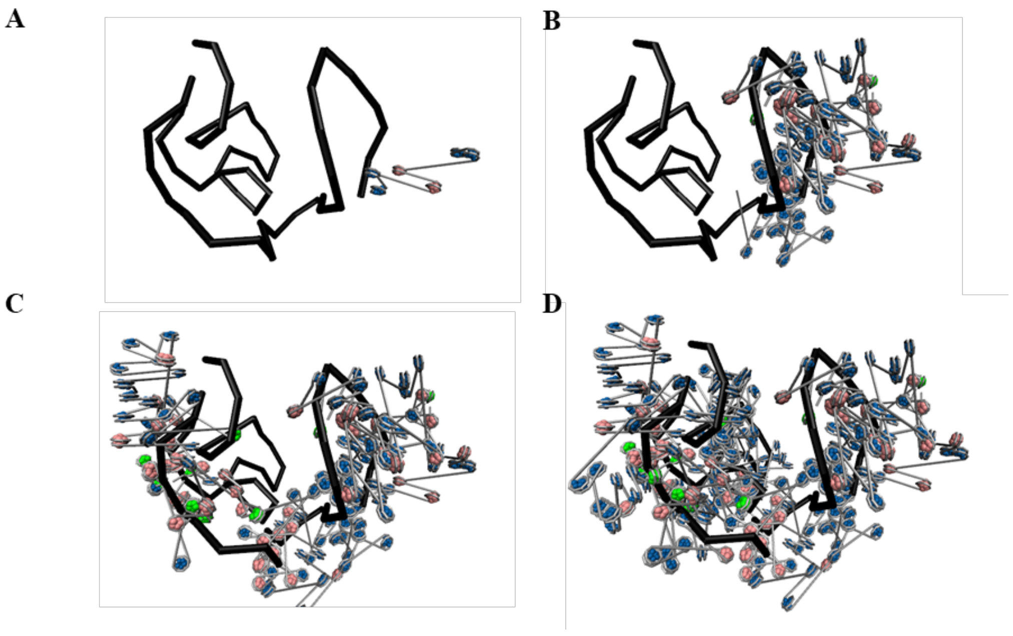


**Figure S4.** Illustration of the back mapping process placing nucleosome resolution chromatin model following the backbone model (black line).

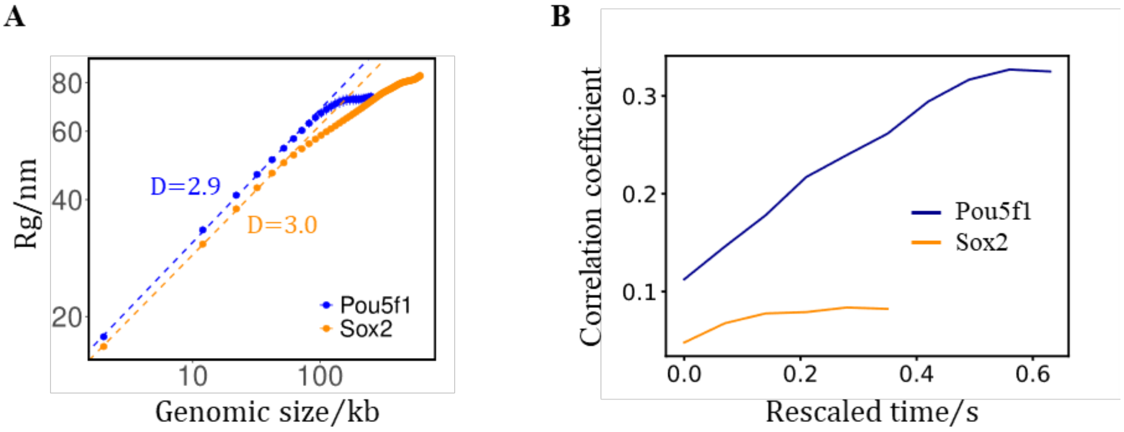


**Figure S5. A.** Radius of gyration – genomic length relations of continuous regions taken from the Sox2 and Pou5f1 loci. The x and y scales are logarithmic. **B.** Mean of Pearson Correlation coefficients between normalized pair-contact frequencies calculated from time $0 \sim \tau$ of any pair of replicas.

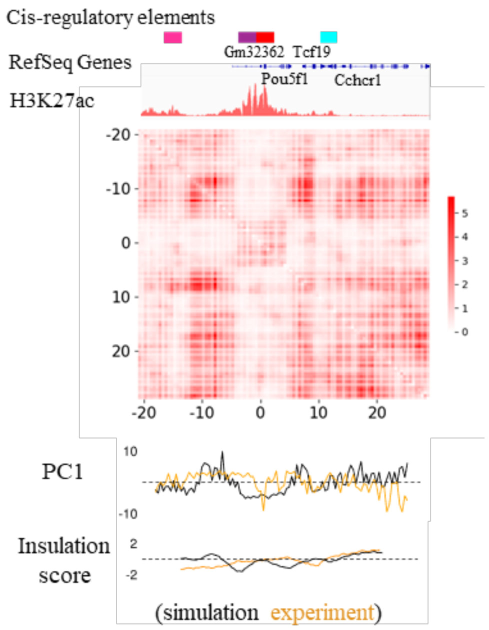


**Figure S6.** PC1 of principal component analysis and insulation score calculated from time $0.14s \sim 0.7s$ from 9 *Pou5f1* locus replicas compared with PC1 and insulation score calculated from experimental contact map.

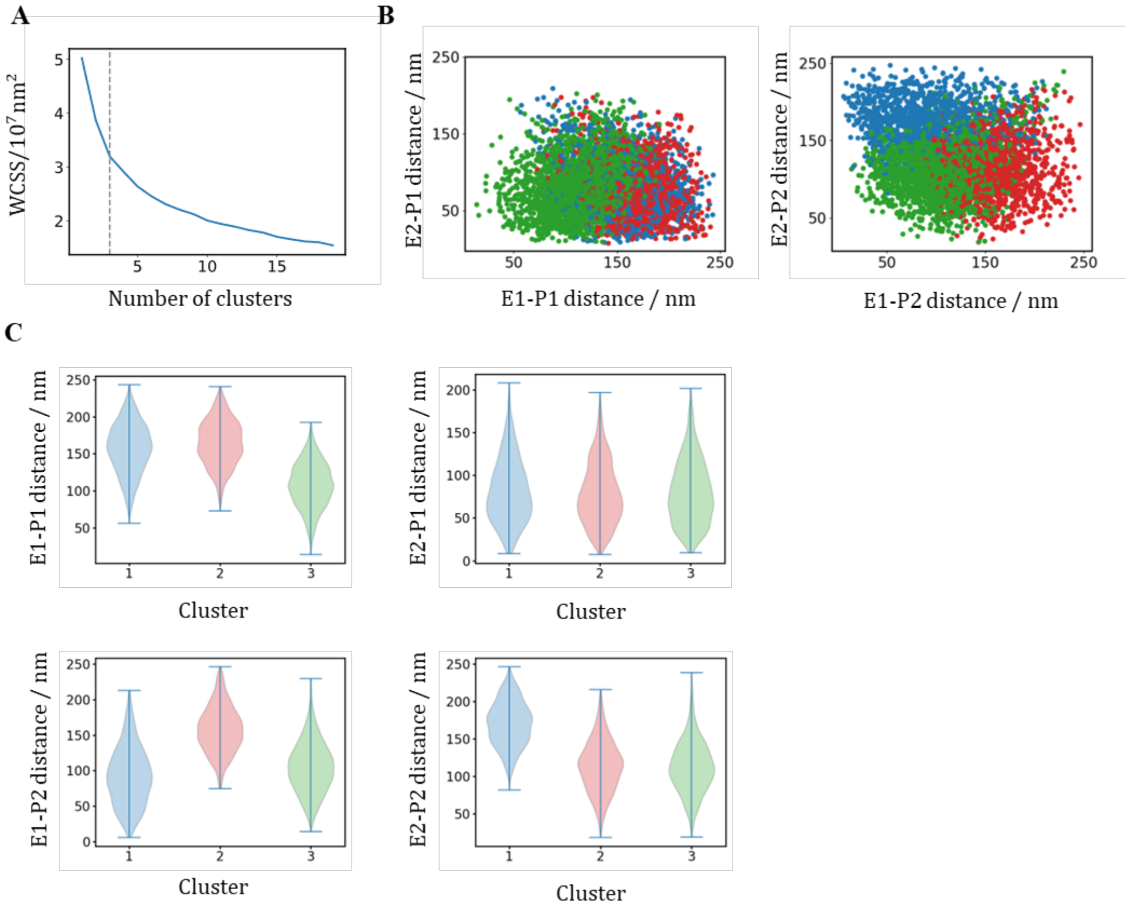


**Figure S7.** K-means clustering of simulated strictures described by regulatory elements pair distances. **A.** Within-cluster-sum-of-squares (WCSS) as a function of number of clusters k used in K-means clustering. Dashed line indicates the chosen “elbow” point (k=3) **B.** Scatter plots of enhancer-promoter distances in 3 clusters. **C.** Violin plots of enhancer-promoter distances in 3 clusters.

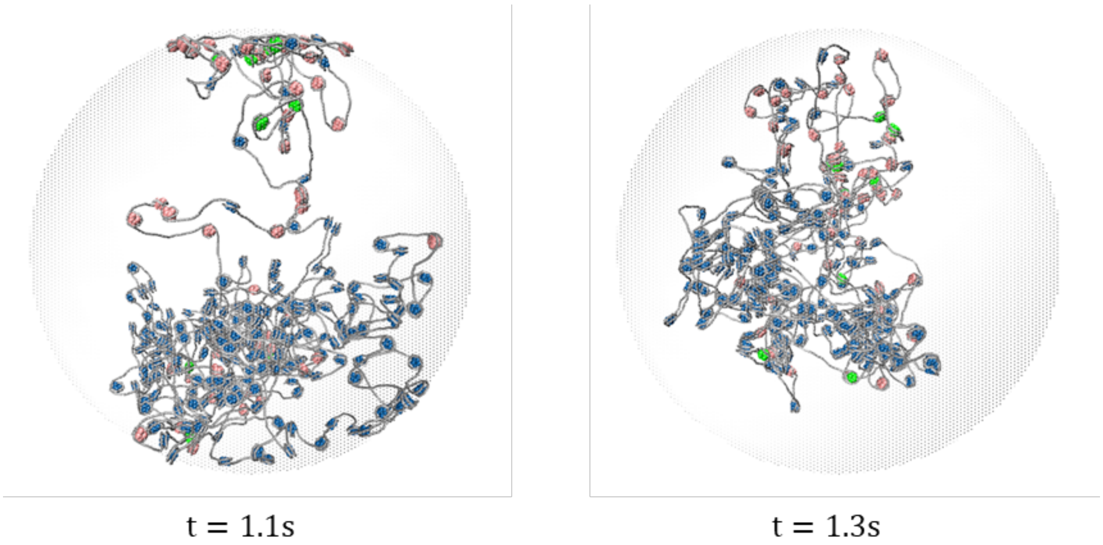


**Figure S8.** Representative structure snapshots from the extended (1.4s) simulation of Pou5f1 locus, with acetylated nucleosomes located on the periphery of the structures.

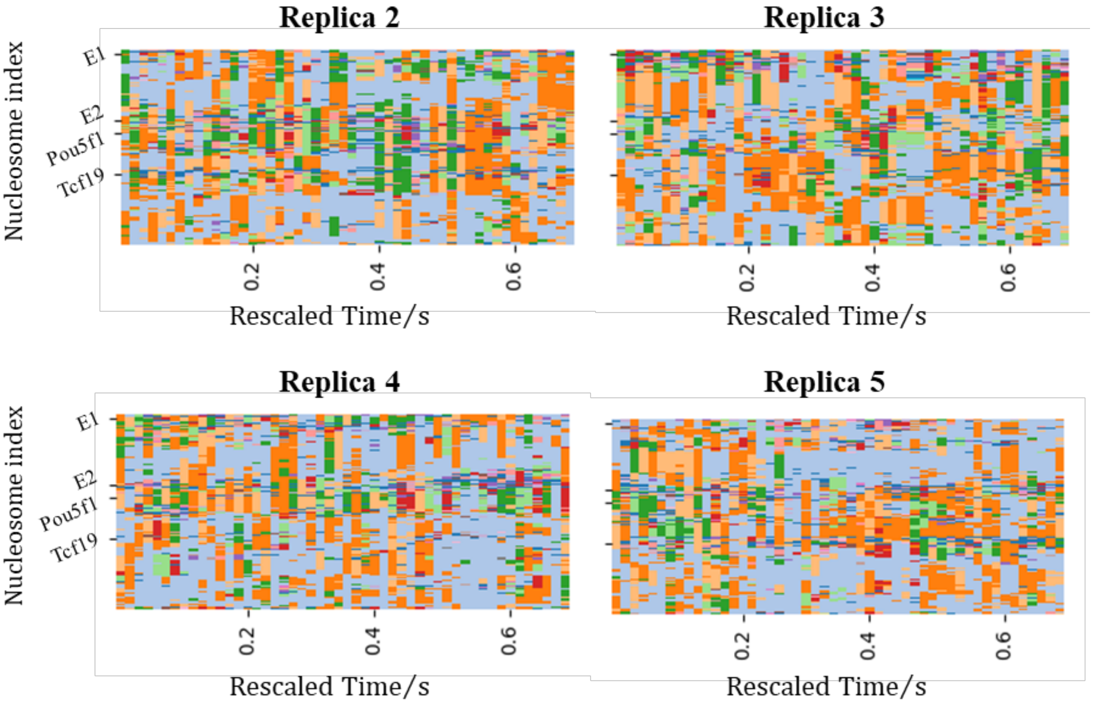


**Figure S9.** Example time series of domains identified from the $0.7s$ Pou5f1 locus simulation trajectories. Domains are indicated by different colors. Calculated every 0.014s.
